# Structural and evolutionary insights into FcαμR recognition of polymeric immunoglobulins

**DOI:** 10.64898/2026.11.03.764125

**Authors:** Mai Zhang, Ruixue Zhang, Yuxin Wang, Junyu Xiao

**Author notes:** Corresponding author: (R.Z.); (J.X.). These authors contributed equally to this work (Mai Zhang, Ruixue Zhang).

## Abstract

Polymeric immunoglobulins (pIgs) are central to adaptive immunity in jawed vertebrates. FcαμR is a bispecific pIg receptor that recognizes both IgM and IgA, but the molecular basis has been unclear. Here we determine cryo-EM structures of mammalian FcαμR bound to IgM or IgA and show that FcαμR engages each ligand in a 1:1 stoichiometry, with higher affinity for IgM. The FcαμR-binding site overlaps with those of pIgR, and also with the preferred FcμR-binding site on IgM, yet distinct features distinguish FcαμR from both pIgR and FcμR. Functionally, FcαμR efficiently internalizes both IgM and IgA through a pathway independent of the di-leucine motif. Structures of amphibian pIgR and FcαμR bound to IgM further shed light on the evolutionary diversification of the FcαμR/pIgR family. Together, these findings define the structural basis of FcαμR-mediated pIg recognition and provide a framework for understanding the evolution of pIg receptors.

## Introduction

Polymeric immunoglobulins (pIgs) are evolutionarily conserved across jawed vertebrates and contribute importantly to immune defense^1^. In mammals, pIgs include IgM and IgA^2^. IgM is an evolutionarily early antibody class, and the initial responder in humoral immunity; it circulates predominantly as a joining chain (J-chain) associated pentamer (pIgM), though J-chain independent polymers, particular hexamer, also occur. IgA is the most abundant antibody at mucosal surfaces and typically forms a J-chain linked dimer (dIgA), with tetrameric and pentameric forms present at lower abundance^3^. The J-chain is essential for the assembly of pIgM and dIgA and also their mucosal secretion, a process mediated by the polymeric immunoglobulin receptor (pIgR)^4,5^. The extracellular portion of pIgR, known as the secretory component (SC), remains bound to mucosal IgM and IgA, forming secretory IgM (SIgM) and secretory IgA (SIgA) complexes^4,6–10^.

Antibodies mediate their effector functions by engaging specific Fc receptors expressed on immune cells^11^. In mammals, IgM and IgA antibodies are recognized by several Fc receptors, including the bispecific receptors FcαμR and pIgR, the IgM-specific receptor FcμR, and the IgA-specific receptor FcαRI, among others^12^. Notably, FcαμR, together with pIgR and FcμR, are thought to share a common evolutionary origin and belong to the same PIGR family^13^.

FcαμR, also known as CD351, was first identified and named for its bispecificity for IgM and IgA^14^. FcαμR is primarily expressed on marginal zone (MZ) B cells, follicular dendritic cells (FDCs), and macrophages^14–16^, playing roles in both humoral immunity and inflammatory responses. For example, it influences germinal center formation and IgG3 affinity maturation by regulating the endocytosis and retention of immune complexes (ICs) in response to T cell-independent (TI) antigens^14–19^. Furthermore, it is involved in regulating the production of pro-inflammatory cytokines by MZ B cells upon lipopolysaccharide stimulation^20,21^. FcαμR is also expressed in several non-lymphoid tissues, including the liver, kidney, intestine, testis and placenta, where it plays roles in maintaining local immune homeostasis^22–24^. Notably, the kidney shows the highest transcript levels of FcαμR in both humans and mice^22^, and there are indications for its implication in IgA nephropathy (IgAN). Under physiological conditions, FcαμR is expressed at low levels in glomerular mesangial cells; nevertheless, upon deposition of galactose-deficient (Gd-) IgA1-ICs, local inflammation-derived IL-1 upregulates FcαμR expression^25^. This upregulation, together with other mesangial receptors such as CD71, appears to further promote ICs deposition^26^.

The molecular mechanism underlying the FcαμR–pIgs interaction remains incompletely understood. Recently, an early homolog of FcαμR was identified in *Xenopus laevis*, which displays a conserved tissue-expression profile and dual binding specificity for two pIgs isotypes, IgM and IgX^13,27^. These findings prompted us to investigate the mechanisms of pIgs recognition by FcαμR in both mammals and amphibians, and to perform cross-species comparisons alongside analyses against other related receptors. Together, these insights shed light on the function and evolution of pIgs and the PIGR family.

## Results

### Cryo-EM structures of FcαμR–Fcμ–J and FcαμR–Fcα–J

The extracellular domain (ECD) of mammalian FcαμR consists of a compact immunoglobulin-like (Ig-like) D1 domain and a long, disordered stalk region containing multiple O-glycosylation sites. Since the stalk of the related receptor FcμR is implicated in facilitating multivalent binding^28,29^, we utilized the entire FcαμR-ECD to study its interaction with IgM and IgA. We generated C-terminally biotinylated human FcαμR-ECD, and then immobilized it on a streptavidin surface plasmon resonance (SPR) sensor chip to mimic membrane presentation. The core region of pIgM (IgM-Fc pentamer with J-chain, Fcμ–J) and dIgA1 (IgA1-Fc dimer with J-chain, Fcα–J) were used as analytes in the SPR experiments. FcαμR binds Fcμ–J with higher affinity (apparent K_D_ = 73.6 ± 2.1 nM; Fig. 1a) than Fcα–J (apparent K_D_ = 7.5 ± 4.2 µM; Fig. 1b). The interaction with Fcα–J was characterized by fast association and dissociation kinetics, whereas binding to Fcμ–J displayed slower kinetics.

**Fig. 1.**
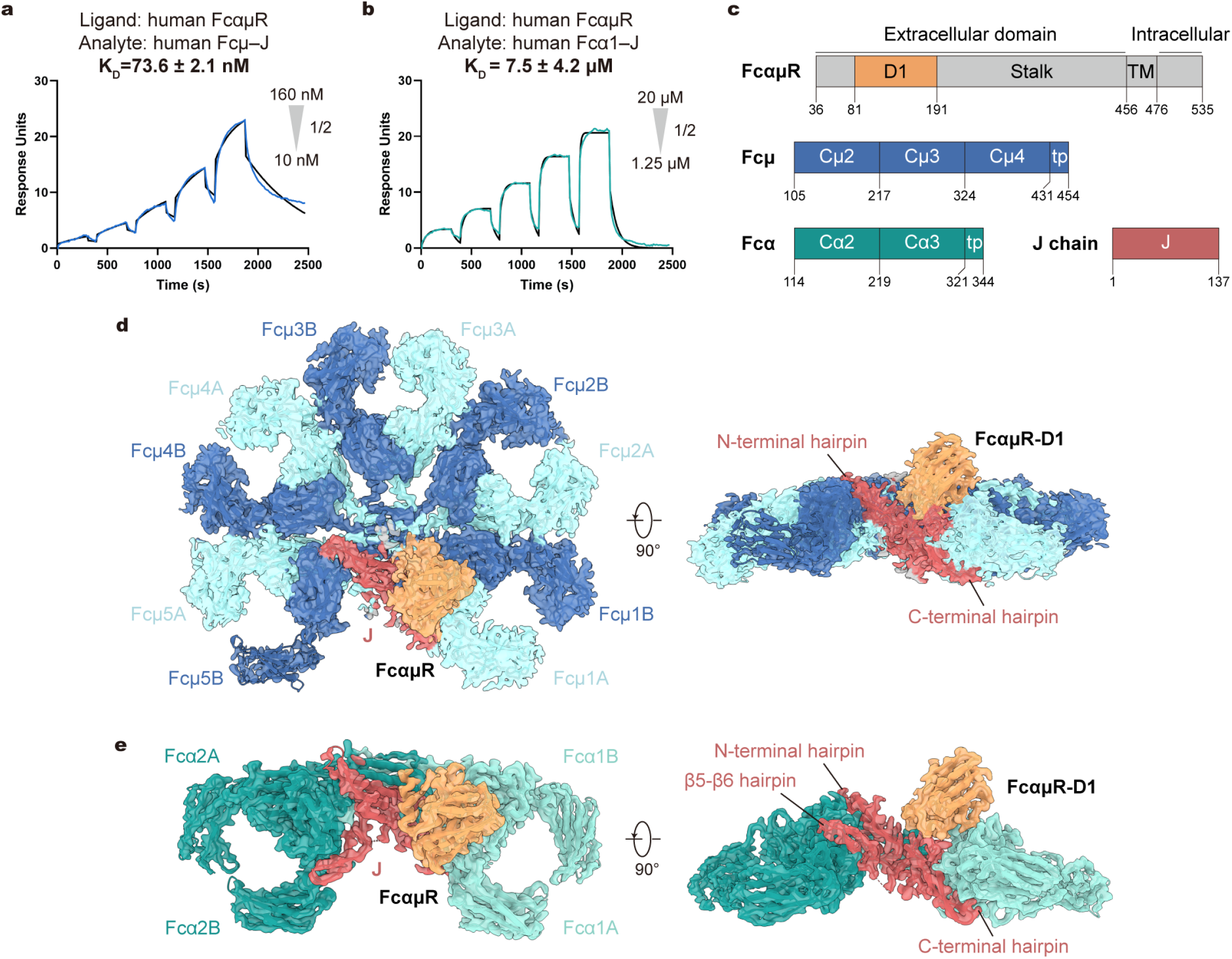
Cryo-EM structures of the FcαμR–Fcμ–J and FcαμR–Fcα–J complexes. **a–b**, SPR analyses of human FcαμR interactions with Fcμ–J (**a**) or Fcα–J (**b**). Two-fold serial dilutions of purified Fcμ–J (from 160 nM to 10 nM) or Fcα1–J (from 20 μM to 1.25 μM) were injected over C-terminal immobilized FcαμR-ECD, using a single-cycle kinetics program. Experimental sensorgrams (colored lines) are overlaid with fitted binding curves (black lines). The reported affinity values are the mean of three independent replicates, presented in Extended Data Fig. 1. **c,** Domain organizations of mouse FcαμR, Fcμ, Fcα and J chain. TM, transmembrane; tp, tailpiece. **d–e,** Cryo-EM reconstructions of mouse FcαμR–Fcμ–J (**d**) and FcαμR–Fcα–J (**e**) complexes shown in two views. FcαμR-D1 and J-chain are shown in sandy brown and red. The Fcμ molecules are shown in two shades of blue, while Fcα1 and Fcα2 are shown in two shades of cyan. Similar color schemes are used in all figures unless otherwise indicated.

For structural analyses, we succeeded in obtaining mouse FcαμR-ECD in complex with either mouse Fcμ–J or Fcα–J by co-expression in HEK293F cells, and then performed cryo-electron microscopy (cryo-EM) analyses (Fig. 1c; Extended Data Figs. 2 and 3; Table 1). Cryo-EM structures of the FcαμR–Fcμ–J and FcαμR–Fcα–J complexes were determined at global resolutions of 2.93 Å and 2.49 Å, respectively (Fig. 1d, e). In both structures, only a single FcαμR molecule engages with one Fcμ–J or Fcα–J complex, forming a 1:1 stoichiometry. We did not observe any particle classes containing more than one FcαμR molecule bound to one Fcμ–J or Fcα–J. This finding stands in sharp contrast to the FcμR–Fcμ– J interaction, which frequently involves multiple FcμR molecules binding to Fcμ–J in a multivalent fashion^28,29^. The single FcαμR molecule binds at a site nearly identical to those occupied by the D1 domain of pIgR-SC in SIgM/SIgA^6–10^, and by the R1 site of FcμR in the FcμR–Fcμ–J complex^28^. Although the samples contained the full-length FcαμR-ECD, only weak densities were observed for the stalk region, indicating inherent flexibility.

**Table 1.** Cryo-EM data collection, refinement and validation statistics.

|  | **mouse**<br>**FcαμR–Fcμ–J**<br>PDB 25GO<br>EMD-80089 | **mouse**<br>**FcαμR–Fcα–J**<br>PDB 25GQ<br>EMD-80091 | ***Xenopus***<br>**pIgR–Fcμ–J**<br>PDB 25OF<br>EMD-80245 | ***Xenopus***<br>**FcαμR–Fcμ–J**<br>PDB 25OG<br>EMD-80246 |
|---|---|---|---|---|
| **Data collection and processing** |  |  |  |  |
| Magnification | 165,000 | 165,000 | 165,000 | 165,000 |
| Voltage (kV) | 300 | 300 | 300 | 300 |
| Electron exposure (e<sup>-</sup>/Å<sup>2</sup>) | 60 | 60 | 60 | 60 |
| Defocus range (μm) | -1.3 to -1.7 | -1.3 to -1.7 | -1.0 to -1.4 | -1.0 to -1.4 |
| Pixel size (Å) | 0.74 | 0.74 | 0.74 | 0.74 |
| Initial particle images (no.) | 6,508,991 | 5,705,065 | 4,991,948 | 3,023,859 |
| Final particle images (no.) | 442,157 | 395,844 | 508,376 | 563,788 |
| Symmetry imposed | C1 | C1 | C1 | C1 |
| Map resolution (Å)<br>FSC threshold: 0.143 | 2.93 | 2.49 | 2.76 | 3.15 |
| Map sharpening *B* factor (Å<sup>2</sup>) | -67.3 | -76.2 | -81.5 | -109.6 |
| **Refinement** |  |  |  |  |
| Initial model used (PDB code) | 6KXS, Alphafold3 | 7JG1, Alphafold3 | Alphafold3 | Alphafold3 |
| Model composition |  |  |  |  |
| Non-hydrogen atoms | 19,506 | 9,097 | 23,003 | 21,392 |
| Protein residues | 2,479 | 1,174 | 2,921 | 2,712 |
| Ligands | 11 | 9 | 0 | 0 |
| *B* factors (Å<sup>2</sup>) |  |  |  |  |
| Protein | 78.82 | 18.91 | 53.31 | 72.56 |
| Ligand | 69.21 | 34.97 | / | / |
| R.m.s. deviations |  |  |  |  |
| Bond lengths (Å) | 0.005 | 0.005 | 0.005 | 0.008 |
| Bond angles (°) | 1.094 | 1.026 | 0.709 | 0.965 |
| Validation |  |  |  |  |
| MolProbity score | 1.89 | 1.53 | 1.77 | 1.80 |
| Clashscore | 11.20 | 5.27 | 9.46 | 7.98 |
| Poor rotamers (%) | 0.31 | 0.19 | 0.00 | 0.00 |
| Ramachandran plot |  |  |  |  |
| Favored (%) | 95.27 | 96.27 | 96.03 | 94.72 |
| Allowed (%) | 4.73 | 3.73 | 3.97 | 5.28 |
| Disallowed (%) | 0.00 | 0.00 | 0.00 | 0.00 |

Notably, approximately 10% of particles in the FcαμR–Fcα–J dataset displayed a 2:2 binding stoichiometry. These particles were evenly distributed across the grid and showed no evidence of denaturation or aggregation (Extended Data Fig. 4a, b). Although limited particle numbers and orientational heterogeneity prevented high-resolution reconstruction (Extended Data Fig. 4c, d), the corresponding low-resolution map allows the fitting of two independent 1:1 FcαμR–Fcμ–J structural models. The two complexes adopt a face-to-face arrangement, with the C-termini of both FcαμR molecules oriented toward the same direction (Extended Data Fig. 4e, f). This configuration could reflect a potential mode for bivalent binding and cell-surface recognition of dIgA by FcαμR.

### Molecular details of FcαμR binding to IgM and IgA

Cryo-EM analysis reveals that FcαμR engages both IgM and IgA primarily through three complementarity-determining region (CDR)-like loops within its D1 domain: CDR1 (Ser108– Arg114), CDR2 (Asn134–Tyr136), and CDR3 (Asp177–Leu182) (Fig. 2a, e). CDR1 primarily contacts the J-chain in both complexes (Fig. 2b, f), reminiscent of the CDR1 loop in pIgR-SC^6–10^. CDR2, along with an adjacent β-strand, interacts with both chains of the Fcμ1 (Fcμ1A/Fcμ1B) or Fcα1 (Fcα1A/Fcα1B) subunit (Fig. 2c, g). CDR3 contacts Fcμ1B/Fcα1B and extends to engage tailpiece residues from the opposing Fcμ5B/Fcα2B subunit (Fig. 2d, h).

**Fig. 2.**
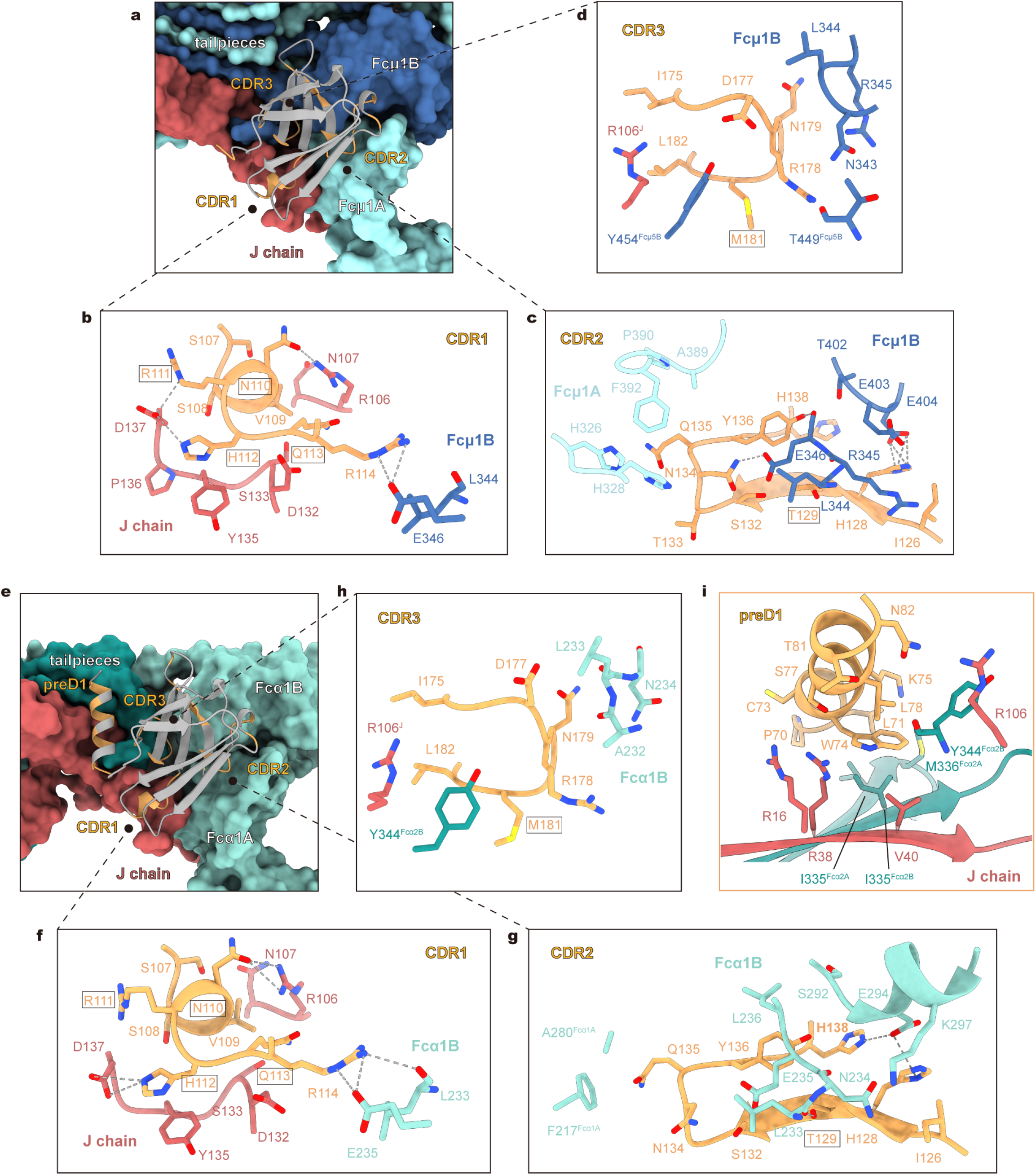
Molecular details of FcαμR interaction with Fcμ–J and Fcα–J. **a**, Binding interface between FcαμR and Fcμ–J. FcαμR-D1 is shown as grey ribbon, with interacting residues highlighted in sandy brown. Fcμ–J is shown as surface. **b–d,** Interactions between FcαμR and Fcμ–J at CDR1 (**b**), CDR2 (**c**) and CDR3 (**d**) regions. Dash lines indicate polar interactions. Boxed residues were mutated in the functional assays presented in Fig. 3. **e,** Binding interface between FcαμR and Fcα–J. FcαμR-D1 is shown as grey ribbon, with interacting residues highlighted in sandy brown. Fcα–J is shown as surface. **f–h,** Interactions between FcαμR and Fcα–J at CDR1 (**f**), CDR2 (**g**) and CDR3 (**h**) regions. Dash lines indicate polar interactions. Boxed residues were mutated in the functional assays presented in Fig. 3. **i,** Interactions between FcαμR and Fcα–J at the preD1 region. The helix mediates hydrophobic and van der Waals interactions with tailpiece residues Ile335^Fcα2A^, Met336^Fcα2A^, Ile335^Fcα2B^, and Tyr344^Fcα2B^, as well as with J-chain residues Arg16^J^, Arg38^J^, Val40^J^ and Arg106^J^.

In the FcαμR–Fcμ–J complex, notable interactions include: within CDR1, Asn110 forms a hydrogen bond with Arg106^J^, Arg111 forms salt bridge with Asp137^J^, and His112 packs against Tyr135^J^ (Fig. 2b). Arg114 additionally contacts Glu346^Fcμ1B^. In CDR2, Asn134 and Tyr136 together interact with Glu346^Fcμ1B^. His128, located on the β-strand N-terminal to CDR2, contacts Glu404^Fcμ1B^ (Fig. 2c). In CDR3, Met181 and Leu182 sandwich Tyr454^Fcμ5B^ (Fig. 2d). In the FcαμR–Fcα–J complex, CDR1-mediated contacts are largely similar (Fig. 2f), whereas interactions involving CDR2 and CDR3 appear somewhat reduced compared to the Fcμ–J complex (Fig. 2g, h). A distinctive feature of this complex is the ordering of a short segment N-terminal to the D1 domain (Pro70–Gly80, designated the preD1 region), which is disordered in the Fcμ–J complex. In the Fcα–J complex, this region adopts a stable helical conformation (Extended Data Fig. 4g) and forms multiple contacts with tailpiece residues from Fcα2A and Fcα2B, as well as with the J-chain (Fig. 2i). Although this preD1 region is present across mammalian species, its sequence is not highly conserved (Extended Data Fig. 4h), and its functional relevance to the interaction between human FcαμR and Fcα–J remains unclear.

The high-resolution FcαμR–Fcα–J map also reveals new features for the Fcα–J core. Each Fcα tailpiece contains a conserved cysteine residue. Consistent with prior studies^8–10^, Cys343 in Fcα1A and Fcα2B forms disulfide bonds with Cys68 and Cys14 of the J-chain, respectively. Our structure further reveals that the tailpieces of Fcα1B and Fcα2A fold back to form an additional inter-chain disulfide bond, with surrounding residues stabilized by hydrophobic packing against adjacent tailpiece chains (Extended Data Fig. 4i).

### FcαμR mutants display reduced binding to IgM and IgA

To assess the functional importance of the interactions identified in our structures, we first generated four FcαμR-ECD mutants and tested their binding by pull-down assays. The N110A/R111A/H112A mutant in CDR1, designed to diminish contact with the J-chain, markedly reduced binding to both Fcμ–J and Fcα–J (Fig. 3a, b). Three additional mutants, Q113N/K115T (CDR1), T129N/V131T (CDR2), and M181N/F183T (CDR3), were all engineered to introduce N-linked glycosylation sites, thereby imposing a steric blockade in each region. Indeed, each of these glycosylation-site mutants completely abolished interaction with both Fcμ–J and Fcα–J (Fig. 3a, b).

**Fig. 3.**
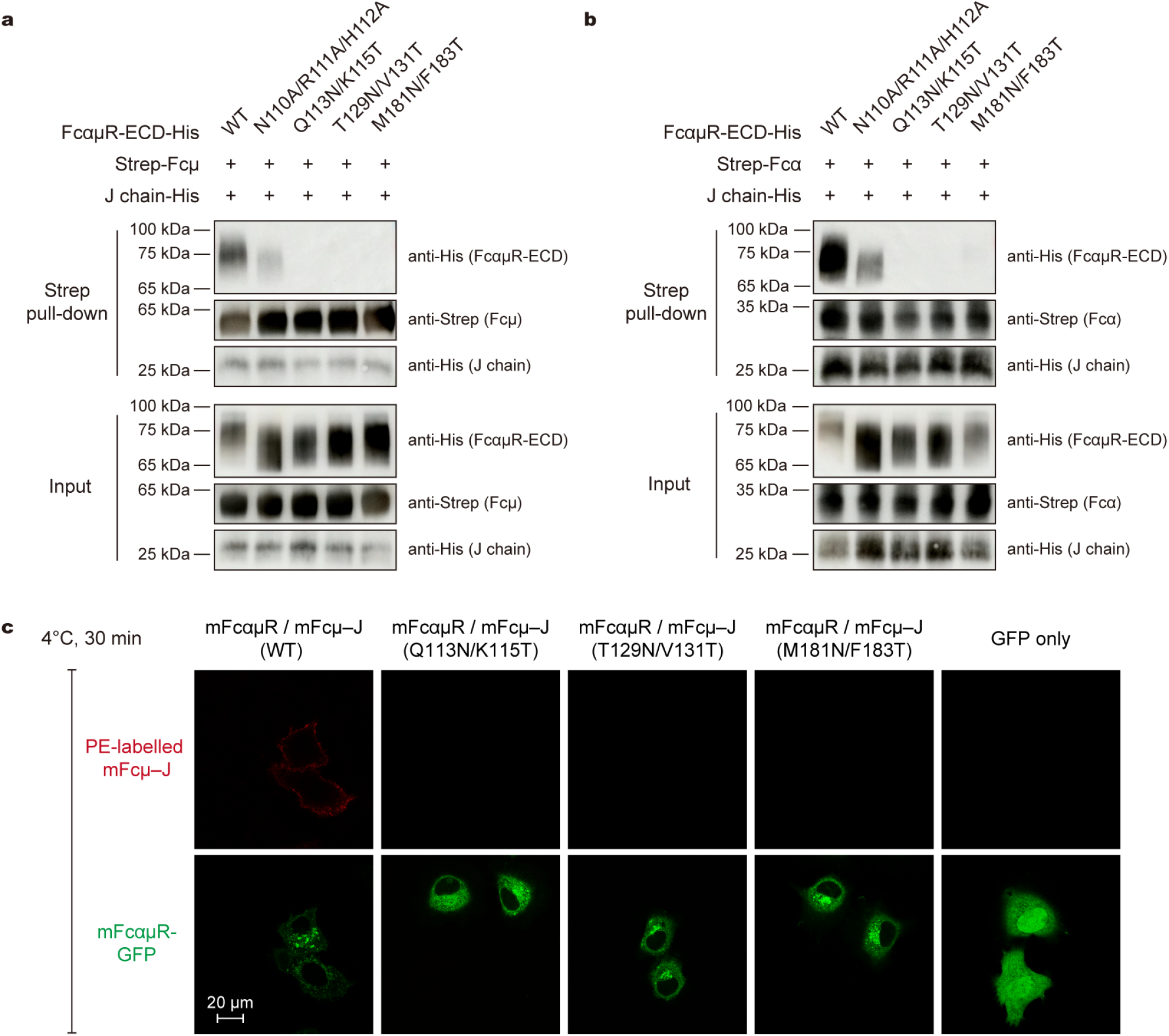
FcαμR mutants displayed reduced interaction with Fcμ–J and Fcα–J. **a–b**, Strep pull-down assay of FcαμR mutants binding to Fcμ–J (**a**) and Fcα–J (**b**). Compared to WT FcαμR, the N110A/R111A/H112A mutant displayed reduced interaction, whereas the Q113N/K115T, T129N/V131T and M181N/F183T mutants displayed abolished interaction. The mutated residues are boxed in Fig. 2. The results were analyzed by SDS-PAGE and detected through immunoblotting using antibodies to the Strep tag and His tag. **c,** Confocal microscopy assay of FcαμR mutants binding to Fcμ–J. HeLa cells expressing WT FcαμR or the indicated glycosylation mutants (Q113N/K115T, T129N/V131T, M181N/F183T) were incubated with twin-strep-tagged Fcμ–J. Binding was detected using PE-conjugated streptavidin (red), while FcαμR was visualized via its C-terminal GFP tag (green). Fcμ–J binding was observed for WT FcαμR (33 out of 40 GFP⁺ cells). In contrast, no Fcμ–J binding was detected in any of the three glycosylation mutants (0 out of 14, 0 out of 15, and 0 out of 14 GFP⁺ cells, respectively) or the GFP-only negative control (0 out of 12 GFP⁺ cells). Images are representative of cells examined at 4°C. See also Fig. 4 for WT FcαμR results.

We further evaluated FcαμR binding in a cellular context by expressing wild-type (WT) or mutant FcαμR as C-terminal GFP fusions on the surface of HeLa cells. Cells expressing any of the three glycosylation mutants showed no detectable binding to soluble Fcμ–J, in contrast to WT receptor (Fig. 3c). Together, these data highlight the essential role of the structural interface we identified and confirm that the observed contacts are critical for FcαμR engagement with IgM and IgA.

### FcαμR mediates internalization of IgM and IgA

Prior studies demonstrated that mouse FcαμR can mediate internalization of IgM-ICs^14^ and human FcαμR can mediate internalization of both IgM and IgA^30,31^. The endocytic activity underpins the receptor’s involvement in humoral immunity^14–17^. To extend these functional insights and to assess the conservation of FcαμR-mediated endocytosis between human and mice, we directly visualized the binding and internalization of IgM and IgA by human and mouse FcαμR using confocal fluorescence microscopy. HeLa cells were transfected with full-length FcαμR bearing a C-terminal GFP tag. Incubation with species-matched Fcμ–J or Fcα–J at 4°C revealed strong binding of both ligands to FcαμR-expressing cells (Fig. 4a). Upon shifting the temperature to 37°C, Fcμ–J and Fcα–J were efficiently internalized within an hour, by both human and mouse FcαμR (Fig. 4b).

**Fig. 4.**
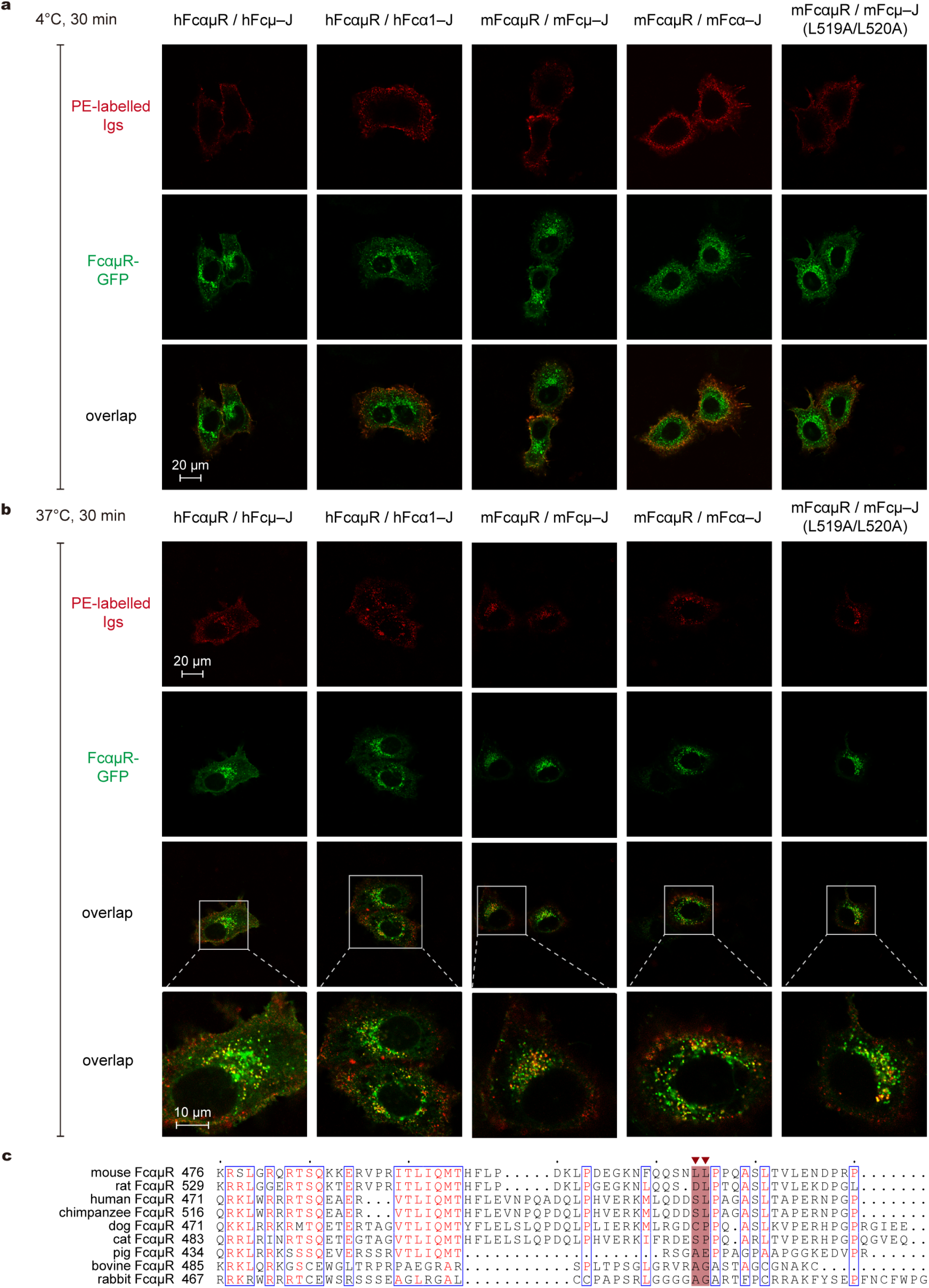
FcαμR mediates internalization of Fcμ–J and Fcα–J. HeLa cells expressing FcαμR were incubated with twin-strep-tagged Fcμ–J or Fcα. The localization of Fcμ–J or Fcα–J was detected using PE-conjugated streptavidin (red), while FcαμR was visualized via its C-terminal GFP tag (green). Experiments were performed twice with consistent results. **a,** Representative images of cells observed at 4°C. For human FcαμR: 22 out of 25 GFP⁺ cells were hFcμ–J⁺; 21 out of 24 GFP⁺ cells were hFcα1–J⁺. For mouse FcαμR: 33 out of 40 GFP⁺ cells were mFcμ–J⁺; 31 out of 35 GFP⁺ cells were mFcα–J⁺; for the L519A/L520A mutant: 24 out of 29 GFP⁺ cells were mFcμ–J⁺. **b,** Representative images of cells observed at 37°C. For human FcαμR: 42 out of 46 GFP⁺ cells were hFcμ–J⁺; 44 out of 49 GFP⁺ cells were hFcα1–J⁺. For mouse FcαμR–GFP: 25 out of 28 GFP⁺ cells were mFcμ–J⁺; 29 out of 32 GFP⁺ cells were mFcα–J⁺; for L519A/L520A: 52 out of 59 GFP⁺ cells were mFcμ–J⁺. **c,** Multiple sequence alignment of the intracellular region in mammalian FcαμR across nine species. The position corresponding to the di-leucine motif in mouse is highlighted in red.

Notably, mutation of the di-leucine motif in the intracellular region of mouse FcαμR (L519A/L520A) did not impair its endocytic activity (Fig. 4b). This finding contrasts with earlier reports^14^; the reason for the discrepancy remains unclear. Nevertheless, the result is consistent with the poor conservation of this motif across mammals (Fig. 4c) and with the observation that human FcαμR, which lacks the motif, still mediates robust internalization. Together, these data indicate that FcαμR employs an endocytic mechanism that can operate independently of the di-leucine motif.

### Comparison of pIgR, FcαμR, and FcμR

The three members of the PIGR family, pIgR, FcαμR, and FcμR (TOSO/FAIM3), are encoded within a conserved syntenic region of the mammalian genome (e.g., human chromosome 1q32)^14^. While all three receptors engage pIgs primarily through a membrane-distal immunoglobulin-like D1 domain, their binding specificities differ^32^. pIgR is a dual-specificity receptor that transports both IgM and IgA across mucosal epithelia. FcαμR also binds both isotypes. In contrast, FcμR is generally considered an IgM-specific receptor^33^.

Indeed, the binding mode of FcαμR closely resembles that of the D1 domain of pIgR (Fig. 5a–d), consistent with their high sequence homology and similar CDR-like loop lengths (Fig. 5i); whereas FcμR exhibits structural differences and consequential binding preferences. Although FcμR primarily engages the same canonical site (the R1 site) as pIgR and FcαμR, it can readily bind other Fcμ units within pIgM or SIgM, leading to multivalent interactions (Fig. 5a)^28,29^. FcαμR, however, exhibits a 1:1 engagement to pIgM. This constraint is sterically imposed by its extended CDR1 loop, which is four residues longer than that of FcμR and is optimally accommodated at the R1 site through contacts with the J-chain (Fig. 5e, j). Structural modeling shows that if FcαμR were to bind at alternative sites occupied by FcμR, its longer CDR1 would clash either with a neighboring Fcμ subunit (at the R2–R4 or R2′–R4′ sites; Fig. 5f) or with the β5–β6 region of the J-chain (at the R1′ site; Fig. 5g). Thus, FcαμR prefers to form a monovalent interaction with the IgM pentamer. Conversely, the longer CDR3 loop of FcμR penetrates more deeply into the IgM binding pocket than its FcαμR counterpart (Fig. 5e), contributing to FcμR’s higher affinity and selectivity for IgM. Further functional specialization arises from differential regulation by the circulating factor CD5L (AIM). While CD5L does not affect FcμR binding^34,35^, it potently inhibits the FcαμR–IgM interaction^34^. Modeling suggests this inhibition results from a steric clash between the FcαμR CDR1 loop and the SRCR3 domain of CD5L (Fig. 5h), suggesting FcαμR preferentially engages CD5L-less IgM.

**Fig. 5.**
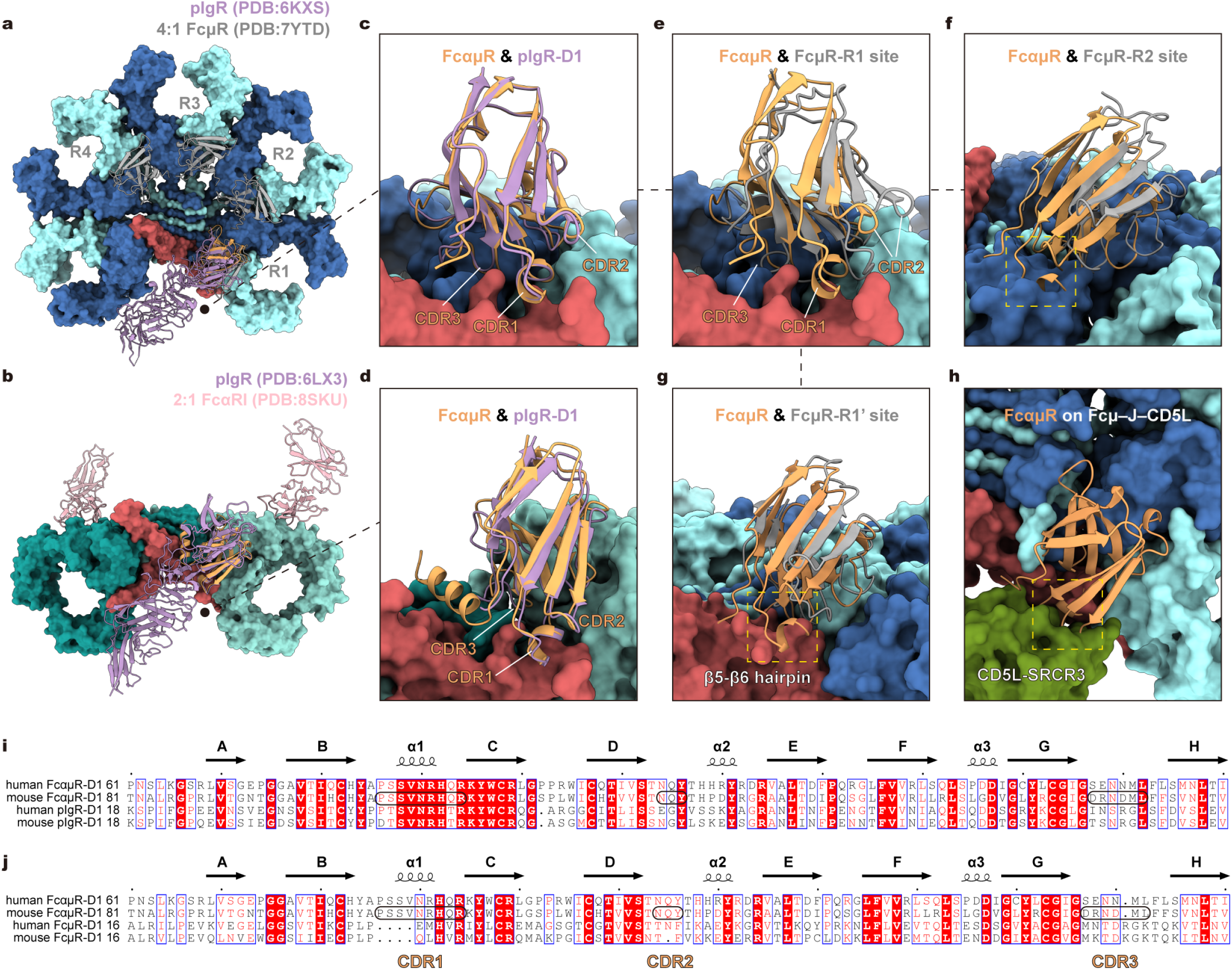
IgM and IgA recognition by FcαμR compared with other receptors. **a**, Binding positions of FcαμR, pIgR (PDB: 6KXS) and FcμR (R1–R4 sites; PDB: 7YTD) on Fcμ–J. FcαμR, pIgR and FcμR are shown in sandy brown, plum and gray, respectively. Similar color schemes are used in all figures unless otherwise indicated. **b,** Binding positions of FcαμR, pIgR (PDB: 6LX3) and FcαRI (PDB: 8SKU) on Fcα–J. FcαμR, pIgR and FcαRI are shown in sandy brown, plum and pink, respectively. **c,** Comparison of FcαμR-D1 binding sites with pIgR-D1 on Fcμ–J. **d,** Comparison of FcαμR-D1 binding sites with pIgR-D1 on Fcα–J. **e–g,** Comparison of FcαμR-D1 binding sites with FcμR-D1 at R1 (**e**), R2 (**f**) or R1’ (**g**) sites on Fcμ–J. Clear steric clashes are observed between FcαμR and the Fcμ–J plane at the R2 site (PDB: 7YTD), and between FcαμR and the J chain at the R1′ site (PDB: 7YSG), as indicated by yellow boxes. These clashes preclude the multivalent binding of Fcμ–J by FcαμR. **h,** Superposition of FcαμR-D1 onto the Fcμ–J–CD5L complex (PDB: 8WYR). CD5L is shown in olive green. Clash between FcαμR-D1 and the SRCR3 domain of CD5L is highlighted with a yellow box. **i–j,** Sequence alignments between FcαμR-D1 and pIgR-D1 (**i**) or FcμR-D1 (**j**) in human and mouse. The three CDR regions are highlighted with rectangles in the mouse FcαμR sequence.

### Evolutionary divergence of PIGR family receptors in binding to pIgs

The African clawed frog (*Xenopus laevis*) provides a valuable model for investigating the early diversification of immunoglobulins (Igs) and their receptors. *Xenopus* possesses two pIg isotypes: IgM and IgX. IgX is primarily secreted across mucosal surfaces and is considered a functional analogue of mammalian IgA, and assembles as a J-chain-independent hexamer in vivo (Fig. 6a)^36,37^. A recently identified *Xenopus* homolog of FcαμR (xlFcαμR) binds both IgM and IgX^13^. Comparing to the *Xenopus* pIgR (xlpIgR), which contains four Ig-like domains (D1, D2, D4, D5) and lacks the D3 domain present in mammalian pIgR, xlFcαμR comprises only three domains (D1, D2, D4; Fig. 6a).

**Fig. 6.**
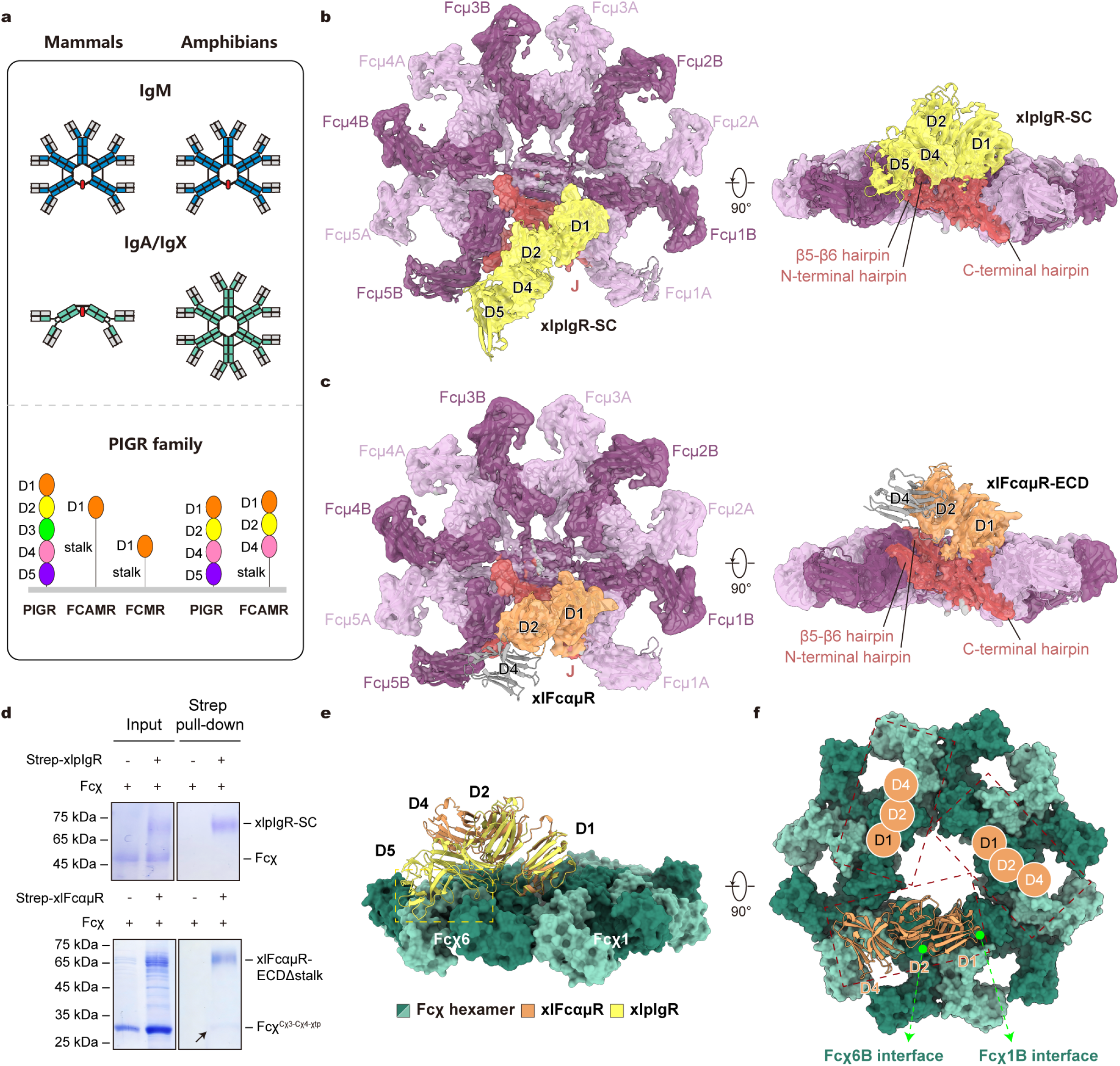
Cryo-EM structures of the xlpIgR–Fcμ–J and xlFcαμR–Fcμ–J complexes. **a**, Polymeric immunoglobulins and PIGR family members in mammals and amphibians, depicting the domain organization of IgM and IgA/IgX, as well as pIgR, FcαμR, and FcμR. **b,** Cryo-EM reconstruction of the xlpIgR–Fcμ–J complex shown in two views. The Fcμ molecules are shown in two shades of purple, while J-chain is shown in red. xlpIgR-ECD is shown in yellow. Similar color schemes are used in all figures unless otherwise indicated. **c,** Cryo-EM reconstruction of the xlFcαμR–Fcμ–J complex shown in two views. xlFcαμR is shown in sandy brown. The positions of the D2 and D4 domains were assigned based on the cryo-EM density map and by fitting an AlphaFold3-predicted model, respectively. **d,** Strep pull-down assay assessing receptor–Fcχ hexamer interactions. The xlpIgR-ECD shows no detectable binding to Fcχ (top panel). In contrast, xlFcαμR-ECDΔstalk binds the Fcχ^cχ3-cχ4-χtp^ hexamer, as indicated by the black arrow (bottom panel). **e,** Superposition of xlpIgR-ECD and xlFcαμR-ECD onto the Fcχ hexamer (PDB: 9UO6). Matches were performed between Fcμ1B and Fcχ1B. The Fcχ molecules are colored in two shades of green. xlpIgR-ECD and xlFcαμR-ECD are shown as yellow and sandy brown cartoons. Clash between xlpIgR-D5 and Fcχ6 is highlighted with a yellow box. **f,** Three potential xlFcαμR-binding sites are located on the same side of the Fcχ hexamer. The interfaces formed by xlFcαμR-D1 are indicated with lime arrows. See also Extended Data Fig. 8k for an enlarged view of the interface.

To compare these early PIGR-family receptors, we purified xlpIgR-ECD or xlFcαμR-ECD in complex with *Xenopus* Fcμ–J (xlFcμ–J) and determined their structures by cryo-EM (Extended Data Figs. 6 and 7; Table 1). The xlpIgR–Fcμ–J and xlFcαμR–Fcμ–J complexes were resolved at resolutions of 2.76 Å and 3.15 Å, respectively (Fig. 6b, c). The overall architecture of xlFcμ–J resembles its human counterpart. Both xlpIgR and xlFcαμR exhibit a 1:1 binding to xlFcμ–J, and the interactions involve only their D1 domains (Extended Data Fig. 8a–c). In the xlpIgR–Fcμ–J complex, the interaction between xlpIgR and xlFcμ–J is highly similar to that seen between human pIgR and Fcμ–J (Extended Data Fig. 8d–f), despite differences in domain organizations of the two pIgR molecules: the D1, D4, and D5 domains of xlpIgR occupy positions similar to the corresponding domains in human pIgR, whereas its D2 domain aligns with human D3 (Fig. 6b)^6,7^. In the xlFcαμR–Fcμ–J complex, the D1–D2 domains adopt a conformation similar to those in xlpIgR, whereas the D4 domain is largely disordered (Fig. 6b, c). Compared to mammalian FcαμR, xlFcαμR primarily contacts the J-chain and the Fcμ1B chain; interactions with Fcμ1A are markedly weaker, and no contacts are made with the tailpiece region (Fig. 2a–d and Extended Data Fig. 8g–j).

We next assessed receptor binding to hexameric IgX-Fc (Fcχ) by pull-down assay. Consistent with previous observations^13^, only xlFcαμR was able to bind to Fcχ, whereas xlpIgR showed no detectable interaction (Fig. 6d). Superposition of the xlpIgR–Fcμ–J structure onto the Fcχ hexamer^38^ reveals a steric clash between the D5 domain of xlpIgR and Fcχ, explaining its lack of binding; in contrast, xlFcαμR, which lacks the D5 domain, would experience less clash (Fig. 6e and Extended Data Fig. 8k). Although the CDR1 loop of xlFcαμR-D1 may still pose some steric hindrance with an adjacent Fcχ subunit, similar to the clash observed between mammalian FcαμR and Fcμ, the presence of Gly47 within this loop likely increases its conformational flexibility, thereby mitigating potential clashes and facilitating engagement with Fcχ (Extended Data Fig. 8j). Moreover, the C6 symmetry of the Fcχ hexamer presents multiple equivalent binding sites, enabling avidity effects to compensate for the loss of J-chain-mediated contacts (Fig. 6f).

## Discussion

Our results show how FcαμR engages pIgs, including IgM and IgA in mammals, and IgM and IgX in amphibians. In mammals, FcαμR binds to Fcμ–J and Fcα–J with a 1:1 stoichiometry and exhibit higher affinity for Fcμ–J. Although our affinity and kinetic measurements differ with some prior reports^39,40^, we note that C-terminal immobilization used here offers some advantages over conventional CM5-chip coupling in preserving native receptor orientation.

FcαμR contributes to a range of physiological processes. FcαμR is found primarily on follicular dendritic cells (FDCs), subsets of B cells, and macrophages in the spleen, regulates humoral immune responses through its capacity to internalize and retain ICs^14–17^. The conservation of this function is supported by the presence of *Xenopus* FcαμR on B cells and myeloid-derived antigen-presenting cells (APCs), cell types analogous to mammalian FDCs^27^. However, the precise mechanism governing FcαμR endocytosis remains unclear. In this study, we visualized the endocytic processes of IgM and IgA via human and mouse FcαμR, and found that the di-leucine motif, known to promote FcγRIIB internalization^14^, is dispensable for mouse FcαμR function. Notably, this motif is not conserved in other mammalian species, suggesting the existence of a separate, evolutionarily conserved endocytic mechanism that awaits discovery. Furthermore, evidence suggests that FcαμR may enhance LPS-induced TLR4 oligomerization in an pIgs-independent manner^20,21,41^, highlighting a potential role in innate immunity and inflammation that warrants further investigation. Beyond the spleen, FcαμR is also expressed in several non-lymphoid tissues, where it helps maintain local immune homeostasis at both mucosal and non-mucosal sites^22–24^. Significantly, in IgAN, FcαμR has been implicated as a key mediator that promotes the mesangial deposition of Gd-IgA1-ICs and drives fibrotic responses in PTECs^25,26,42^. This insufficiently elucidated pathological process warrants further investigation.

As the last structurally characterized member of the PIGR family, FcαμR shares architectural features with both pIgR and FcμR^32^. While its D1 domain displays higher sequence homology to pIgR-D1, the overall domain organization of mammalian FcαμR more closely resembles that of FcμR. The evolutionary origins of FcαμR and pIgR remain unresolved. Although pIgR has often been regarded as the earlier-evolving member, several lines of evidence now suggest that FcαμR may represent the primordial receptor of this family. First, in cartilaginous fish, the most ancient jawed vertebrates, polymeric IgM serves as a first-line defense, likely predominant in T-independent responses^13^, a functional niche aligned with FcαμR expression on APCs. Second, although a potential pIgR homolog (containing only two domains) is annotated in teleost fish, its lack of strong homology to mammalian pIgR domains and, more importantly, its unproven transport function cast doubt on whether it represents a *bona fide* pIgR. Third, in amphibians, xlpIgR cannot bind IgX, leaving xlFcαμR as the sole functional receptor for this mucosal isotype. One could think of a plausible evolutionary scenario in which an ancestral FcαμR-like receptor, possessing two or three domains, gave rise to two lineages. In one direction, it acquired additional domains and evolved into pIgR, which later specialized as a mucosal transporter strictly dependent on the J-chain. In mammals, the D5 domain of pIgR further adapted to form a disulfide bond with IgA, enhancing its specificity. In the other direction, the receptor retained only the essential D1 domain and evolved an elongated stalk, descending into mammalian FcαμR and FcμR. The post-amphibian emergence of IgA coincided with J-chain-mediated polymerization and efficient pIgR-facilitated mucosal transport. FcμR, meanwhile, specialized as a high-affinity IgM receptor capable of engaging all forms of IgM.

In summary, here we delineate the molecular mechanism underlying IgM and IgA recognition by the bispecific receptor FcαμR. These findings advance our understanding of the evolution and elusive functions of pIgs and the PIGR receptor family.

## Methods

### Cell culture

HEK293F cells (Thermo Fisher Scientific) were cultured in FreeStyle 293 Expression Medium (Gibco) at 37°C with 5% CO₂ and 55% humidity using a humidified shaker for protein purification. Sf21 and High Five cells (American Type Culture Collection) were maintained using a non-humidified shaker at 27°C in SIM-SF and SIM-HF Medium (Sino Biological), respectively. Hela cells (American Type Culture Collection) were cultured in DMEM (Gibco) supplemented with 1% penicillin-streptomycin (Gibco) and 10% fetal bovine serum (FBS; PAN-Biotech) in a humidified incubator at 37°C with 5% CO2.

### Constructs design

The genes encoding the core regions of mammalian pIgs were subcloned into a modified pcDNA vector containing an N-terminal interleukin-2 (IL-2) signal peptide and a twin-Strep tag. These included: human Fcμ (UniProt P01871, residues 106–453), human Fcα1 (UniProt P01876, residues 123–353), mouse Fcμ (UniProt P01872, residues 104–454), and mouse Fcα (UniProt P01878, residues 114–344). For surface plasmon resonance (SPR) analyses, we also generated constructs without twin-Strep tags. Separately, the genes for full-length human (UniProt P01591) and mouse (UniProt P01592) J-chains (residues 1–159) were cloned into the same vector and engineered with a C-terminal 8×His tag.

Constructs for mammalian FcαμR proteins were tailored for specific applications. For SPR analyses, gene encoding the extracellular domain (ECD) of human FcαμR (UniProt Q8WWV6, residues 17–450) was cloned into the pFastBac vector with an N-terminal melittin signal peptide, a C-terminal 8×His tag, and an Avi tag (sequence: GLNDIFEAQKIEWHE). For cryo-electron microscopy (cryo-EM), mouse FcαμR-ECD (UniProt Q2TB54, residues 36–459) was cloned into the pcDNA vector with an N-terminal IL-2 signal peptide, a C-terminal 8×His tag, and a Flag tag. For pull-down assays, mouse FcαμR-ECD was cloned into the pFastBac vector with an N-terminal melittin signal peptide, a C-terminal 8×His tag and a Flag tag. For confocal microscopy, genes encoding full-length human (residues 1–532) and mouse (residues 1–535) FcαμR were cloned into the pcDNA vector with a C-terminal GFP tag. All FcαμR point mutants were generated using standard PCR-based site-directed mutagenesis.

Constructs for *Xenopus laevis* orthologs were similarly prepared in the pcDNA vector, each containing an N-terminal IL-2 signal peptide. These included: xlFcμ (UniProt Q6GN83, residues 257–605) with an N-terminal Flag tag; Fcχ (UniProt Q6INK3, residues 240–584) and its truncated form Fcχ^Cχ3-Cχ4-χtp^ (residues 360–584), each with an N-terminal 8×His tag; the J chain (UniProt A0A974DYP0, residues 32–164) with a C-terminal 8×His tag; the xlFcαμR-ECD (GenBank OCT94622.1, residues 15–505), xlFcαμR-ECDΔstalk (the stalk-deleted variant, residues 15–355), and xlpIgR-ECD (UniProt A0A974HYK7, residues 19–457), all with a C-terminal twin-Strep tag.

### Protein expression and purification

Since expression of mammalian FcαμR in HEK293F cells alone was inefficient, the baculovirus-insect cell system was utilized. Bacmids for pFastBac constructs (encoding human or mouse FcαμR-ECD) were generated with the Bac-to-Bac system (Invitrogen). Recombinant baculoviruses were produced and amplified in Sf21 insect cells. For protein production, High Five cells at a density of 2 × 10^6^ cells per millilitre were infected with the amplified virus. The conditioned medium was collected 48 hours post-infection by centrifugation and dialyzed into the binding buffer (25 mM HEPES pH 7.4, 150 mM NaCl). FcαμR-ECD proteins were initially purified by Ni-NTA affinity resin (Smart Lifesciences) and eluted with binding buffer containing 500 mM imidazole (Sigma-Aldrich), then further purified by size-exclusion chromatography (SEC) on a Superdex 200 Increase 10/300 GL column (Cytiva) equilibrated with the binding buffer. Mammalian Fcμ–J and Fcα–J were produced in HEK293F cells and purified by Ni-NTA affinity resin followed by SEC, as previously described^6,8^.

To prepare mouse FcαμR–Fcμ–J and FcαμR–Fcα–J complexes for cryo-EM analyses, HEK293F cells were cultured to a density of 1 × 10^6^ cells per millilitre and co-transfected with plasmids encoding mouse FcαμR-ECD, J-chain, and either Fcμ or Fcα at a 1:1:2 ratio using 40 kDa linear polyethylenimine (PEI; Polysciences). For each liter of culture, 1 mg of plasmid mixture was combined with 2 mg of PEI in 50 mL fresh medium, and incubated for 30 min before transfection. Four days post-transfection, the medium supernatant was collected by centrifugation, then concentrated and exchanged into the binding buffer with a Hydrosart Ultrafilter (Sartorius). The FcαμR–Fcμ–J complex was first purified by anti-Flag affinity resin (GenScript) and eluted with the binding buffer supplemented with 0.2 mg mL^-1^ 3×Flag peptide (NJPeptide). The FcαμR–Fcα–J complex was first purified by Strep-Tactin resin (Smart Lifesciences) and eluted with the binding buffer supplemented with 5 mM desthiobiotin (IBA Lifesciences). Both complexes were further purified by SEC on a Superose 6 Increase 10/300 GL column (Cytiva) pre-equilibrated with the binding buffer. Protein fractions were analyzed throughout by SDS-PAGE and visualized with Coomassie staining.

For *Xenopus* proteins, xlFcμ–J was expressed and purified similarly to mammalian Fcμ–J. Fcχ and Fcχ^cχ3-cχ4-χtp^ were expressed and purified as previously described^38^. xlpIgR-ECD, xlFcαμR-ECD, and xlFcαμR-ECDΔstalk were expressed in HEK293F cells and purified by Strep-Tactin resin followed by SEC, as described above. To prepare the xlpIgR–Fcμ–J and xlFcαμR–Fcμ–J complexes for cryo-EM analysis, xlpIgR-ECD or xlFcαμR-ECD were mixed with xlFcμ–J at a 1:2 molar ratio and incubated on ice for 1 h. The complexes were then further purified by SEC and eluted with the binding buffer.

### Surface plasmon resonance

To explore the ability of FcαμR to bind to different IgM and IgA isoforms, human FcαμR with a C-terminal Avi-tag was purified and biotinylated using biotin protein ligase (BirA) in the binding buffer supplemented with 5 mM MgCl_2_, 2 mM ATP, 1 μM BirA, and 0.15 mM D-biotin. BirA was made with reference to the previous protocol^43^. After the reaction, biotinylated protein was further purified by SEC to remove BirA and excess D-biotin.

The SPR experiments were performed on the Biacore T200 (Cytiva), at 25°C with 1×HBS-EP Buffer (Cytiva) containing 10 mM HEPES pH 7.4, 150 mM NaCl, 3 mM EDTA, 0.05% (v/v) surfactant P20. The C-terminal biotinylated FcαμR was immobilized on an streptavidin (SA) chip (Cytiva) according to the manufacturer instructions. The two-fold serial dilutions of purified non-Strep-tagged human Fcμ–J and Fcμ–J proteins were passed over the flow cell at five concentrations using a single-cycle kinetics program. The SPR results were analyzed with Biacore Evaluation Software v2.0.

### Cryo-EM sample preparation and data collection

For the mouse FcαμR–Fcμ–J complex, peak fractions of SEC corresponding to the complex were collected, concentrated to 0.6 mg mL^-1^, and crosslinked with 0.05% glutaraldehyde (Sigma-Aldrich) at 20°C for 10 min. The mouse FcαμR–Fcα–J complex was processed similarly, except that the sample was concentrated to 4.0 mg mL^-1^ and treated with fluorinated octyl maltoside (FOM; Anatrace) at 0.25 mM to improve particle orientations. For the xlpIgR– Fcμ–J and xlFcαμR–Fcμ–J complexes, samples were concentrated to 1.0 mg mL^-1^ and crosslinked under the same conditions.

Cryo-grids were prepared using the Vitrobot Mark IV (FEI). Briefly, a 4 μL aliquot of protein complex was applied to glow-discharged holy-carbon gold grids (Quantifoil, R1.2/1.3, 300-mesh). Grids were blotted for 1.0 s with filter paper and quickly plunged into liquid ethane under 4°C and 100% humidity. Grids were first screened under a 200 kV Talos Arctica microscope equipped with a Ceta camera (FEI). Data collection was carried out on a 300 kV Titan Krios G4 electron microscope equipped with a Falcon 4 camera. Movie stacks were recorded using the EPU software (Thermo Fisher Scientific) at a magnification of 165,000×, corresponding to a calibrated pixel size of 0.74 Å at object scale. The defocus range was set from -1.3 to -1.7 μm for mouse samples, and from -1.0 to -1.4 μm for *Xenopus* samples. Statistics for data collection are summarized in Table 1.

### Cryo-EM data processing, model building and refinement

Workflows for cryo-EM data processing of mouse FcαμR–Fcμ–J and FcαμR–Fcα–J complexes are shown in Extended Data Fig. 2d and 3d, respectively; those for the xlpIgR–Fcμ–J and xlFcαμR–Fcμ–J complexes are provided in Extended Data Fig. 6e and 7e. All datasets were processed using cryoSPARC^44^.

For the mouse FcαμR–Fcμ–J complex, we collected 8,315 movie stacks. Raw movies were motion-corrected using patch motion correction, and patch CTF estimation was performed on dose-weighted micrographs. Micrographs with significant ice contamination or an estimated CTF resolution worse than 5 Å were discarded. Initial particle picking was performed using blob picker. Templates generated from 2D class averages were used for template-based picking and Topaz training, yielding a total set of 6,508,991 particles. After multiple rounds of 2D classification, 801,204 particles were selected, re-extracted, and subjected to ab-initio reconstruction. This was followed by heterogeneous refinement to remove suboptimal particles.

Using the resulting initial model, 541,210 particles underwent homogeneous refinement, yielding a 3.27-Å map. To improve density for the FcαμR region, 3D classification was performed with a focused mask covering FcαμR-D1 and the Fcμ–J core. The final set of 442,157 particles was used for homogeneous refinement, global CTF refinement, and local refinement, resulting in the final map at 2.93-Å resolution.

Data processing for mouse FcαμR–Fcα–J complex followed a similar workflow. From 9,161 micrographs, 5,705,065 particles were auto-picked. Following 2D classification, selected particles were re-extracted and processed through ab-initio reconstruction and heterogeneous refinement. One additional round of heterogeneous refinement was performed using resolution-gradient templates generated from the initial model. The final 395,844 particles containing high-resolution information were then subjected to homogeneous refinement, local CTF refinement, and local refinement, yielding the final map at 2.49-Å resolution.

For the xlpIgR–Fcμ–J complex, 11,228 movie stacks were collected. The auto-picked 4,991,948 particles were processed through several rounds of 2D classification, ab-initio reconstruction, and heterogeneous refinement. The retained particles were used to generate the final 3D reconstruction via homogeneous refinement and non-uniform (NU) refinement, yielding the final map at 2.76-Å resolution. For the xlFcαμR–Fcμ–J complex, 10,012 movie stacks were collected, and 3,023,859 particles were auto-picked. Following an identical processing workflow, the final map at 3.15-Å resolution was obtained.

Model building and refinement were performed as follows. For mouse FcαμR–Fcμ–J complex, Alphafold3-predicted models of mouse IgM-Cμ3-Cμ4 and FcαμR-D1 were docked into the final map using UCSF ChimeraX^45^. Tailpieces were built de novo in Coot^46^, and the J-chain was rebuilt based on the human Fcμ–J structure (PDB: 6KXS). For mouse FcαμR–Fcα– J complex, the mouse Fcα–J structure (PDB: 7JG1) and FcαμR-D1 model were docked. The helix before the N-terminus of FcαμR-D1 was built de novo in Coot. For the *Xenopus* complexes, Alphafold3-predicted models of xlFcμ–J, xlpIgR-ECD, and xlFcαμR-ECD were docked separately, with tailpieces built de novo in Coot. All models were manually adjusted in Coot and automatically refined in Phenix^47^ using real-space refinement with secondary structure and geometry restraints. Local masks for focused classification and refinement were created in UCSF ChimeraX. All reported resolutions are based on the Fourier shell correlation (FSC) 0.143 criterion, with local resolution estimates calculated in cryoSPARC. Structural figures were prepared with UCSF ChimeraX. Amino acid numbering for mouse Fcμ and Fcα and for *Xenopus* Fcμ and Fcχ (UniProt accessions P01872, P01878, Q6GN83, and Q6INK3, respectively) starts from the constant region.

### Pull-down assay

To investigate the interactions between FcαμR variants (wild-type or mutant) and Fcμ–J or Fcα– J, purified proteins were incubated on ice for 1 h. Because Fcμ and Fcα carry an N-terminal twin-Strep tag, the mixture was subsequently incubated with Strep-Tactin resin in the aforementioned binding buffer at 4°C for 1 h with gentle rotation. After washing the resin three times with binding buffer, bound proteins were eluted with buffer supplemented with 5 mM desthiobiotin. Eluates were analyzed by SDS-PAGE and immunoblotting using anti-Strep (1:3,000; A01736-100, GenScript) and anti-His (1:3,000; HT501, TransGen Biotech) antibodies.

To examine the binding of xlpIgR or xlFcαμR to Fcχ, twin-Strep-tagged xlFcαμR-ECDΔstalk or xlpIgR-SC was mixed with Fcχ hexamer and incubated on ice for 1 h. The mixture was then incubated with Strep-Tactin resin at 4°C for 1 h. After three washes with binding buffer, proteins were eluted with 5 mM desthiobiotin and analyzed by SDS-PAGE followed by Coomassie staining.

### Confocal fluorescence microscopy

HeLa cells grown on coverslips were transfected with the indicated full-length FcαμR constructs, each fused with a C-terminal GFP tag, using PEI and cultured for 30 h. The twin-Strep-tagged Fcμ–J or Fcα–J complex was pre-incubated with phycoerythrin (PE)-conjugated streptavidin (1:250 dilution, Invitrogen) at 4°C for 30 min, then added to the cell culture at a final concentration of 15 µg mL^-1^. After further incubation at either 4°C or 37°C for 30 min, cells were washed three times with pre-chilled or pre-warmed PBS, as appropriate, and fixed with 4% paraformaldehyde for 15 min. Imaging was performed on a Zeiss LSM880 confocal microscope using a ×63/1.4 oil objective. All images were processed with Zeiss Zen 3.9 software, with uniform adjustments applied to brightness and contrast.

## Data availability

Cryo-EM density maps and corresponding coordinates for mouse FcαμR–Fcμ–J (EMD-80089; PDB 25GO), mouse FcαμR–Fcα–J (EMD-80091; PDB 25GQ) complexes, *Xenopus* pIgR– Fcμ–J (EMD-80245; PDB 25OF), and *Xenopus* FcαμR–Fcμ–J (EMD-80246; PDB 25OG) complexes, have been deposited in the Electron Microscopy Data Bank and the Protein Data Bank, respectively. Other structural coordinates used in this study are available from PDB with accession code 6KXS, 6LX3, 7JG1, 7YTD, 7YSG, 8SKU, 8WYR, 9UO6. Source data are provided with this paper.

## Acknowledgement

We extend our gratitude to the cryo-EM and high-performance computing platforms at Peking University and Changping Laboratory for assistance with data collection and computation. We thank the National Center for Protein Sciences at Peking University for assistance with the Biacore and confocal microscope facilities. This work received support from the National Natural Science Foundation of China (32325018, 32530053) to J.X., and National Key R&D Program of China (2024YFA1306200) to J.X.

## Author contributions

J.X. conceived and supervised the project. M.Z. performed protein expression and purification for the mouse complexes, and performed cryo-EM data collection, processing, model building and refinement together with Y.W. The analogous experiments for the *Xenopus* complexes were conducted by R.Z. M.Z. carried out structural analyses, pull-down assays, confocal microscopy and SPR experiments. The manuscript was drafted by M.Z. and revised by J.X.

## Additional information

Correspondence and requests for materials should be addressed to Junyu Xiao.

## ED Figure legends

**Extended Data Fig. 1.**
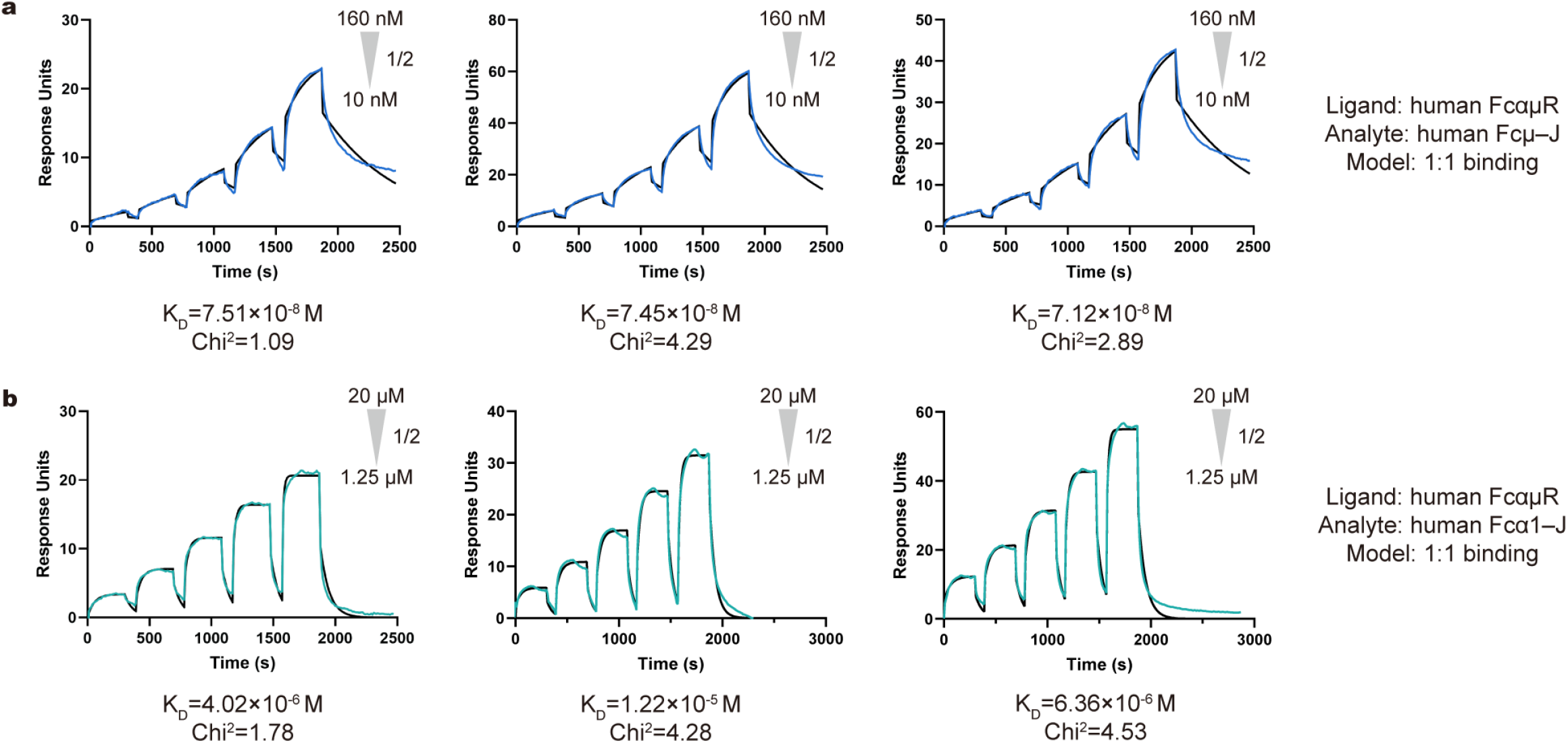
SPR analyses of human FcαμR interactions with IgM and IgA. The SPR experiments were performed in triplicate by passing two-fold serial dilutions of purified Fcμ–J (**a**) and Fcα–J (**b**) over C-terminal immobilized FcαμR-ECD using a single-cycle kinetics program. The highest and lowest concentrations for each analyte are indicated on the graphs. Experimental sensorgrams (colored lines) are overlaid with fitted binding curves (black lines); corresponding fitting statistics are provided. Binding affinities were determined as follows: Fcμ–J, K_D_ = 73.6 ± 2.1 nM (**a**); Fcα–J, K_D_ = 7.5 ± 4.2 μM (**b**). SPR analyses of human FcαμR interactions with Fcμ–J (**a**) or Fcα–J (**b**).

**Extended Data Fig. 2.**
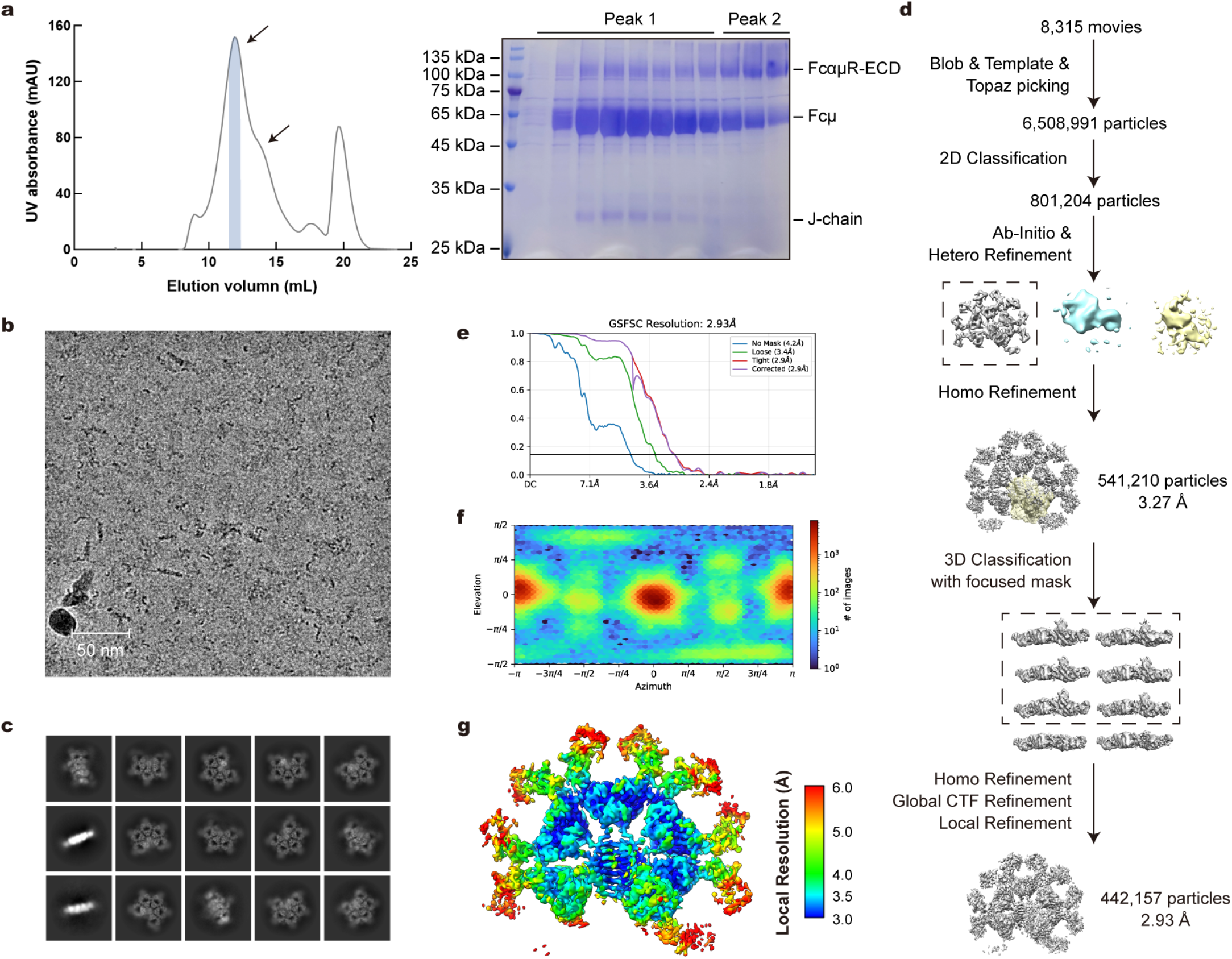
Purification and cryo-EM analysis of the FcαμR–Fcμ–J complex. **a**, Size-exclusion chromatography of the FcαμR–Fcμ–J complex on a Superose 6 Increase column (left) and corresponding SDS-PAGE analysis stained with Coomassie blue (right). Peak 1 corresponds to the FcαμR–Fcμ–J complex, Peak 2 to the excess FcαμR-ECD proteins. Fractions marked with blue shadows were used for cryo-EM analysis. All the purification experiments were repeated at least twice, with similar results. **b,** Representative cryo-EM micrograph of the FcαμR–Fcμ–J complex. Scale bar: 50 nm. **c,** 2D class averages of the FcαμR–Fcμ–J complex. **d,** Workflow of cryo-EM data processing for the FcαμR–Fcμ–J complex. **e,** Gold standard Fourier shell correlation (GSFSC) curves of the final density map. **f,** Angular particle distribution heat map in the final round of local refinement. **g,** Local resolution estimation of the final density map.

**Extended Data Fig. 3.**
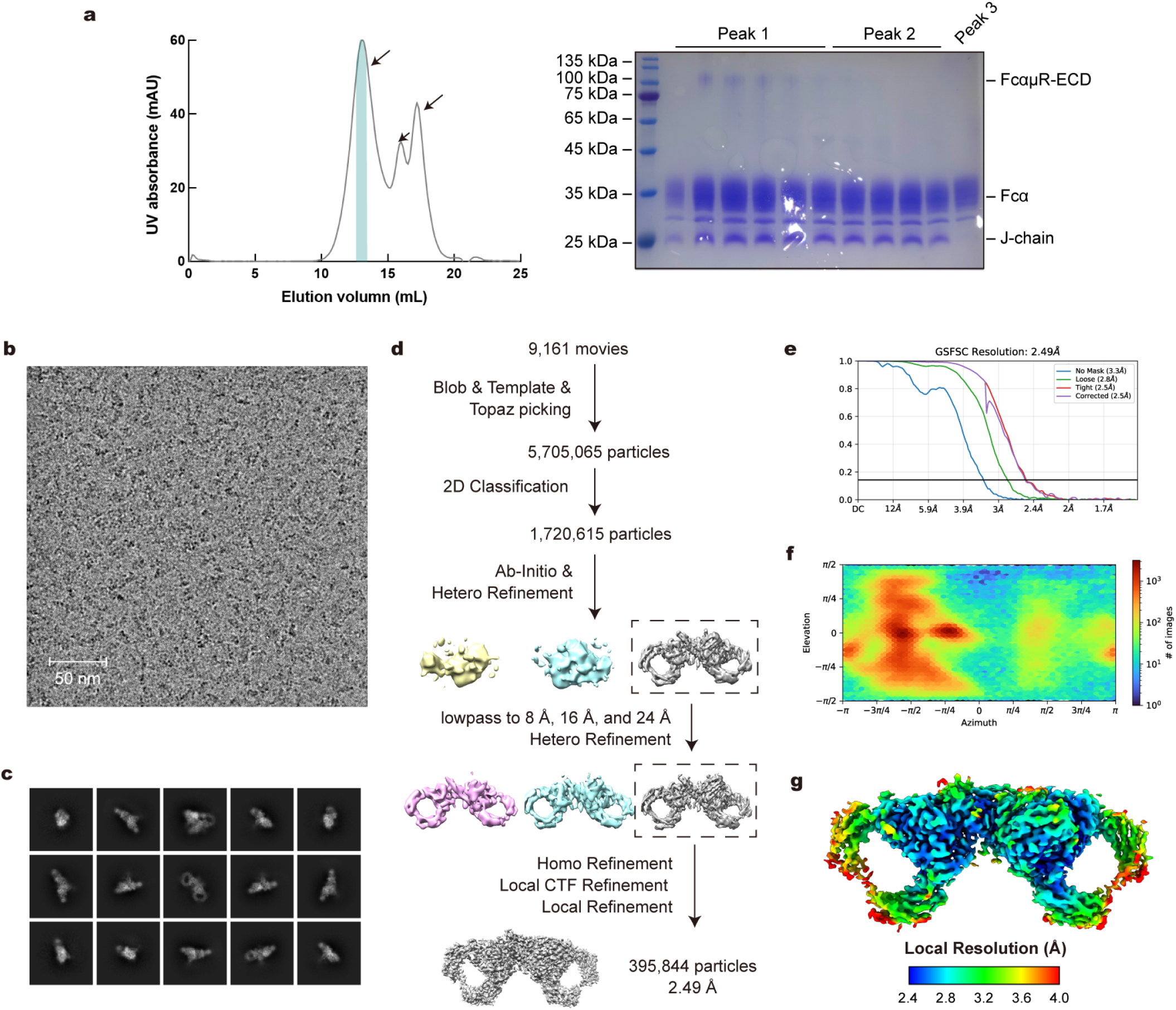
Purification and cryo-EM analysis of the FcαμR–Fcα–J complex. **a**, Size-exclusion chromatography of the FcαμR–Fcα–J complex on a Superose 6 Increase column (left) and corresponding SDS-PAGE analysis stained with Coomassie blue (right). Peak 1 corresponds to the FcαμR–Fcα–J complex, Peak 2 to the Fcα–J complex, and Peak 3 to the Fcα monomer. Fractions marked in dark cyan were used for cryo-EM analysis. **b,** Representative cryo-EM micrograph of the FcαμR–Fcα–J complex. Scale bar: 50 nm. **c,** 2D class averages of the FcαμR–Fcα–J complex. **d,** Workflow of cryo-EM data processing for the FcαμR–Fcα–J complex. **e,** Gold standard Fourier shell correlation (GSFSC) curves of the final density map. **f,** Angular particle distribution heat map in the final round of local refinement. **g,** Local resolution estimation of the final density map.

**Extended Data Fig. 4.**
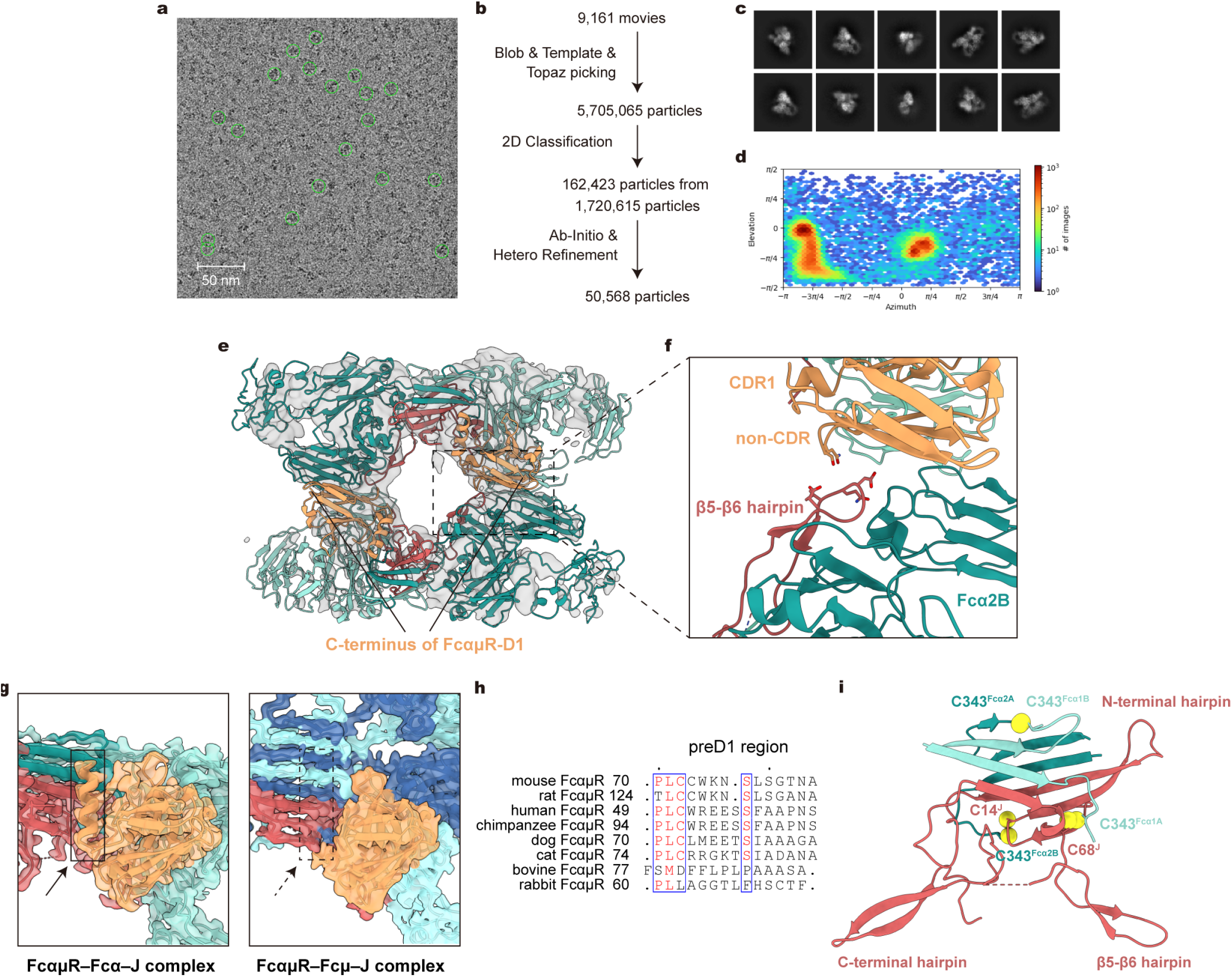
Unique features of the FcαμR–Fcα–J complex. **a**, Cryo-EM micrograph with particles exhibiting a 2:2 binding ratio circled in green. Particles are evenly distributed across the grid with no visible aggregation or signs of denaturation. **b,** Workflow of cryo-EM data processing for the potential 2:2 FcαμR–Fcα–J complex. Approximately 10% of particles (162,423 out of 1,720,615) in the dataset exhibited a 2:2 stoichiometry after 2D classification. See also Extended Data Fig. 3d for the 1:1 complex. **c,** 2D class averages of the potential 2:2 FcαμR–Fcα–J complex. **d,** Angular particle distribution heat map in the final round of refinement job. **e,** Structural analysis of the putative 2:2 FcαμR–Fcα–J complex. The cryo-EM map is shown as a transparent gray surface, with two 1:1 structural models fitted as colored ribbons. The C-termini of both FcαμR molecules face the same side, suggesting a bivalent binding mode for Fcα–J. **f,** Close-up view of the second interface, with key interacting residues shown as sticks. Each FcαμR forms an additional interface where residues from the non-CDR-like loop contact the β5-β6 hairpin of the opposing J-chain and may also interact with the opposite Fcα2B. **g,** Disordered residues at the N-terminal of FcαμR-D1 are induced to form a helical structure when binding to Fcα–J rather than Fcμ–J. **h,** Multiple sequence alignment of the preD1 region in mammalian FcαμR across nine species: human, chimpanzee, mouse, rat, bovine, pig, dog, cat, and rabbit. Corresponding UniProt accession numbers are provided in the same order: Q8WWV6, A0A2I3SL69, Q2TB54, A0A8I6AQN7, A0A3Q1LPX3, K7GP71, A0A8I3MHM2, A0ABI7W8F4, A0A5F9C7C4. **i,** Overall structure of the J-chain and tailpiece residues in the FcαμR–Fcα–J complex. Beyond previous researches^8–10^, we found that the tailpiece ends of Fcα1B and Fcα2A fold back to form a pair of disulfide bond. Tailpieces and J-chain are shown as ribbon, while side chains of cystine residues are represented as yellow spheres.

**Extended Data Fig. 5.**
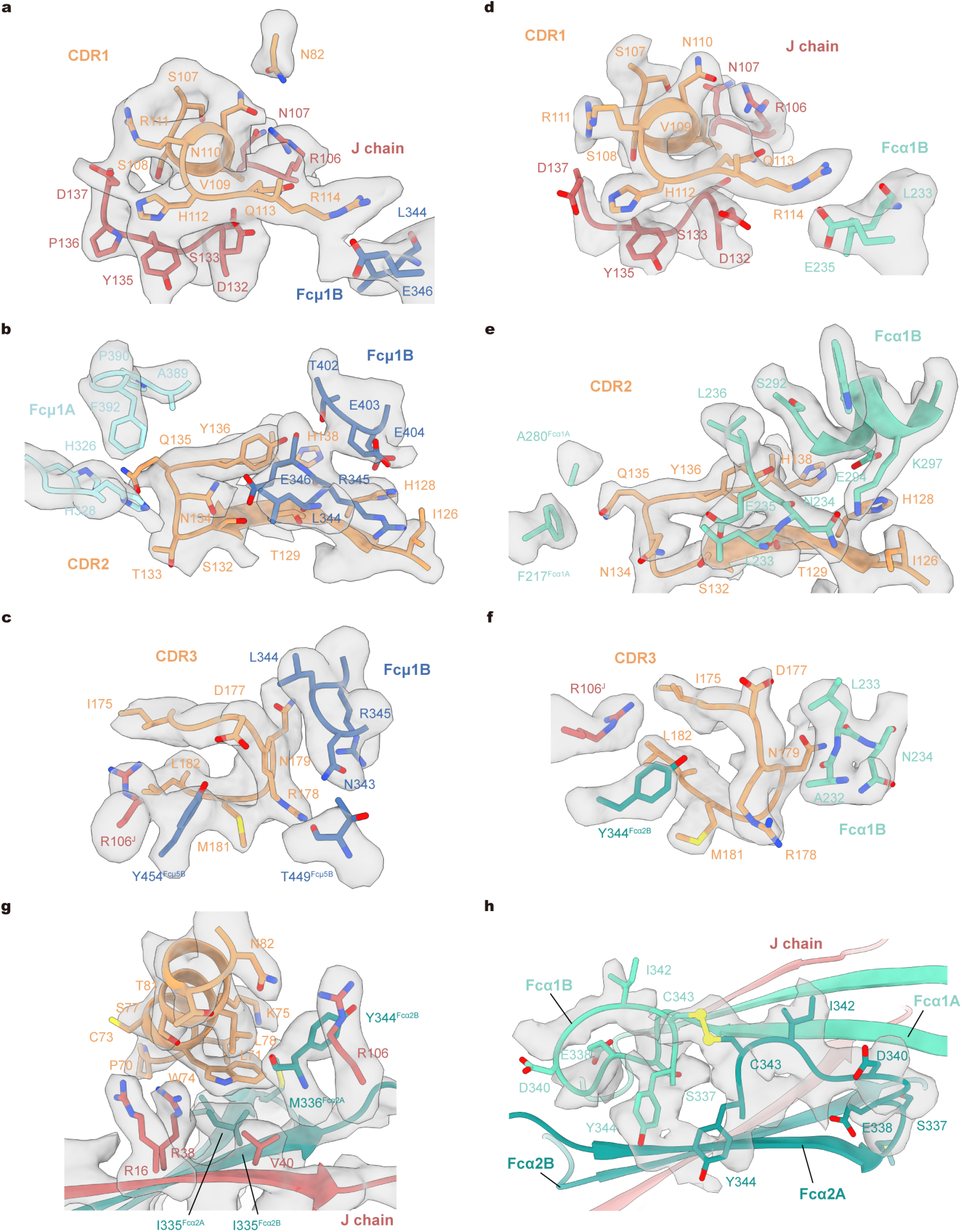
Representative local density maps of the FcαμR–Fcμ–J and FcαμR–Fcα–J complexes. **a–c**, Local density maps of the FcαμR–Fcμ–J interface at CDR regions, corresponding to Fig. 2b–d, respectively. **d–f,** Local density map of the FcαμR–Fcα–J interface at CDR regions, corresponding to Fig. 2f–h, respectively. **g,** Local density maps of the FcαμR–Fcα–J interface at the preD1 helix, corresponding to Fig. 2i. **h,** Local density map of the residues at the end of Fcα1B and Fcα2A, corresponding to Extended Data Fig. 4i.

**Extended Data Fig. 6.**
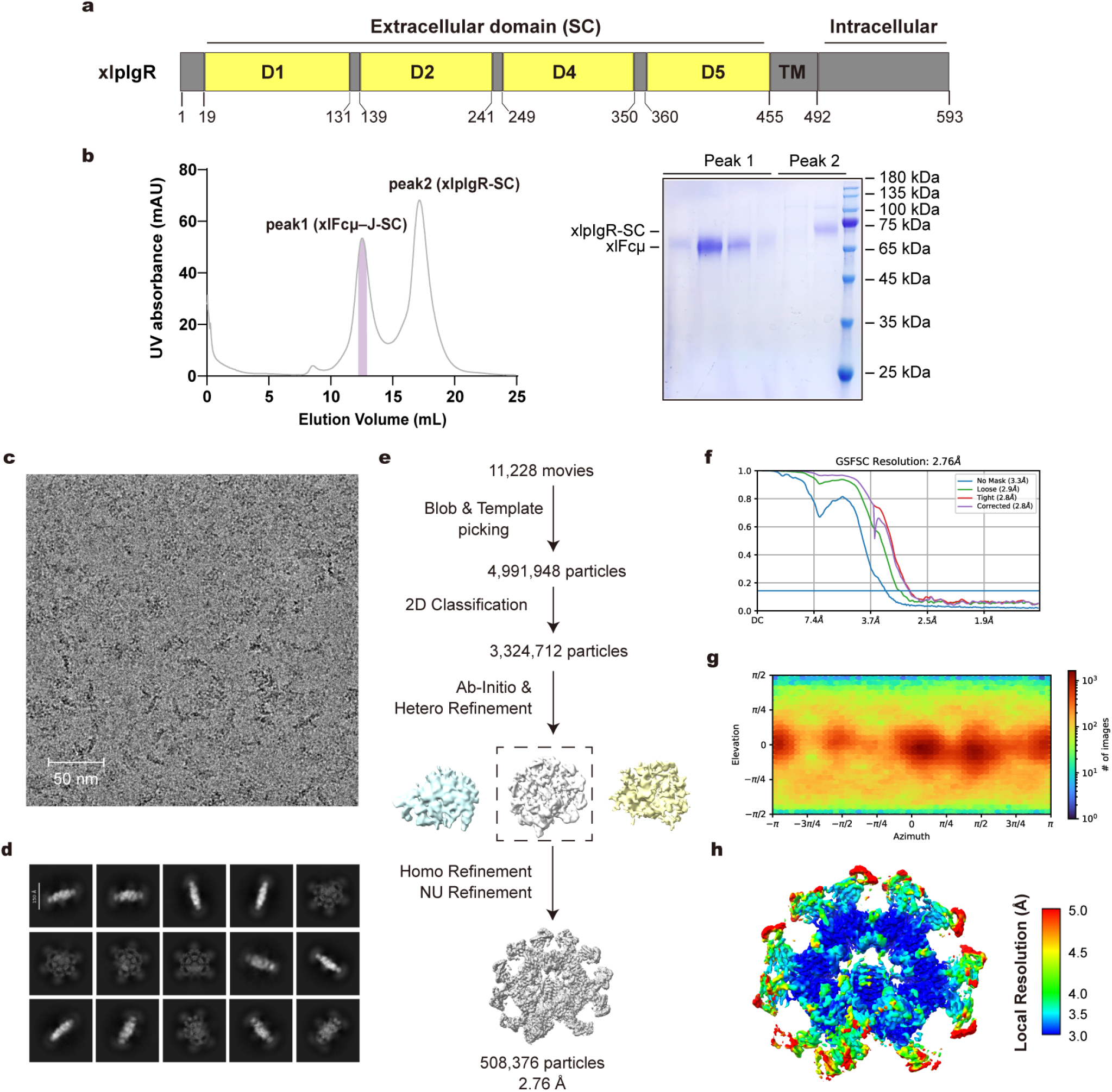
Purification and cryo-EM analysis of the xlpIgR–Fcμ–J complex. **a**, Domain organizations of xlpIgR. TM, transmembrane. **b,** Size-exclusion chromatography of the xlpIgR–Fcμ–J complex on a Superose 6 Increase column (left) and corresponding SDS-PAGE analysis stained with Coomassie blue (right). Peak 1 corresponds to the xlpIgR–Fcμ–J complex, Peak 2 to the excess xlpIgR-ECD proteins. Fractions marked with purple shadows were used for cryo-EM analysis. **c,** Representative cryo-EM micrograph of the xlpIgR–Fcμ–J complex. Scale bar: 50 nm. **d,** 2D class averages of the xlpIgR–Fcμ–J complex. **e,** Workflow of cryo-EM data processing for the xlpIgR–Fcμ–J complex. **f,** Gold standard Fourier shell correlation (GSFSC) curves of the final density map. **g,** Angular particle distribution heat map in the final round of local refinement. **h,** Local resolution estimation of the final density map.

**Extended Data Fig. 7.**
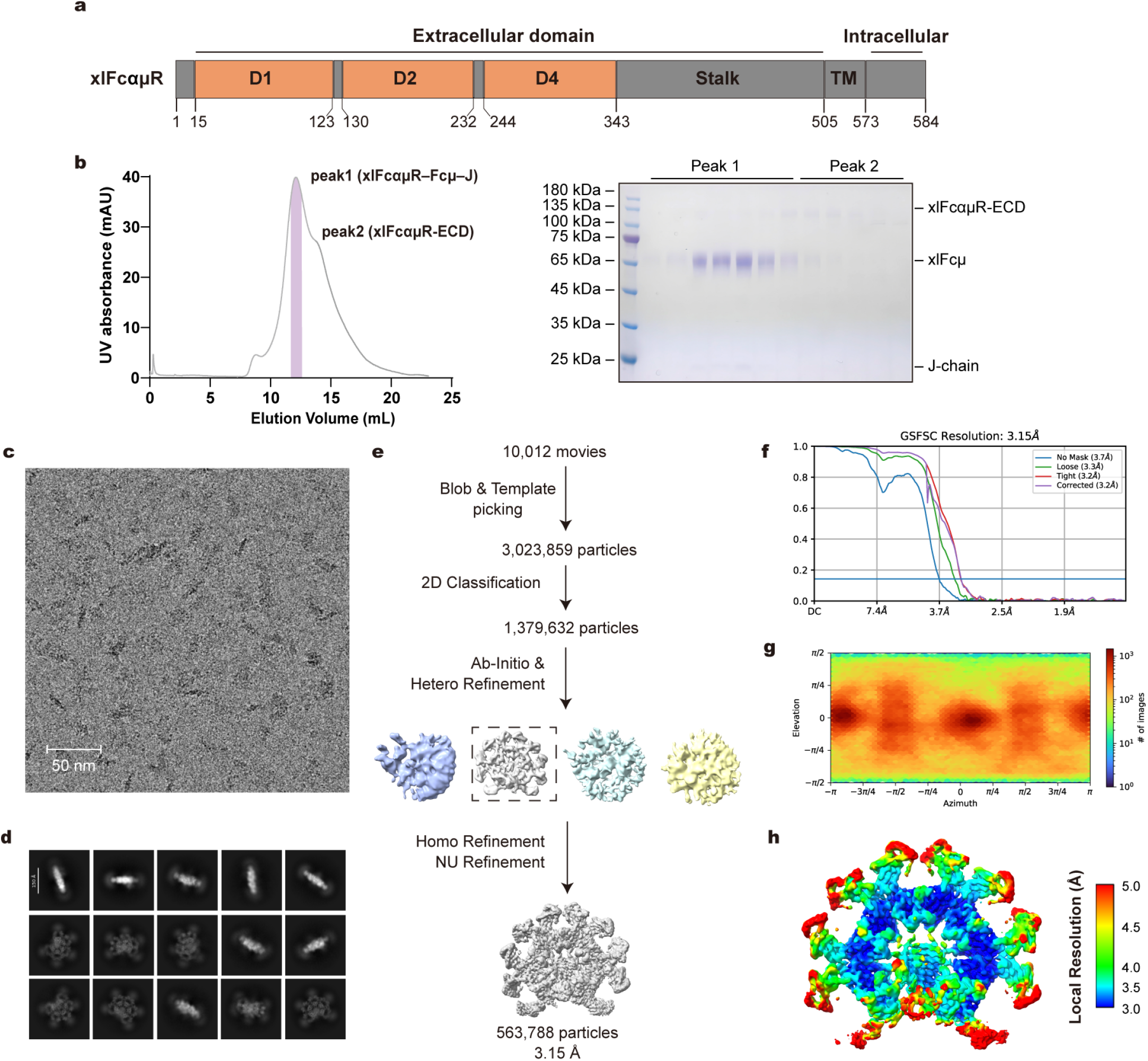
Purification and cryo-EM analysis of the xlFcαμR–Fcμ–J complex. **a**, Domain organizations of xlFcαμR. TM, transmembrane. **b,** Size-exclusion chromatography of the xlFcαμR–Fcμ–J complex on a Superose 6 Increase column (left) and corresponding SDS-PAGE analysis stained with Coomassie blue (right). Peak 1 corresponds to the xlFcαμR–Fcμ–J complex, Peak 2 to the excess xlFcαμR-ECD proteins. Fractions marked with purple shadows were used for cryo-EM analysis. **c,** Representative cryo-EM micrograph of the xlFcαμR–Fcμ–J complex. Scale bar: 50 nm. **d,** 2D class averages of the xlFcαμR–Fcμ–J complex. **e,** Workflow of cryo-EM data processing for the xlFcαμR–Fcμ–J complex. **f,** Gold standard Fourier shell correlation (GSFSC) curves of the final density map. **g,** Angular particle distribution heat map in the final round of local refinement. **h,** Local resolution estimation of the final density map.

**Extended Data Fig. 8.**
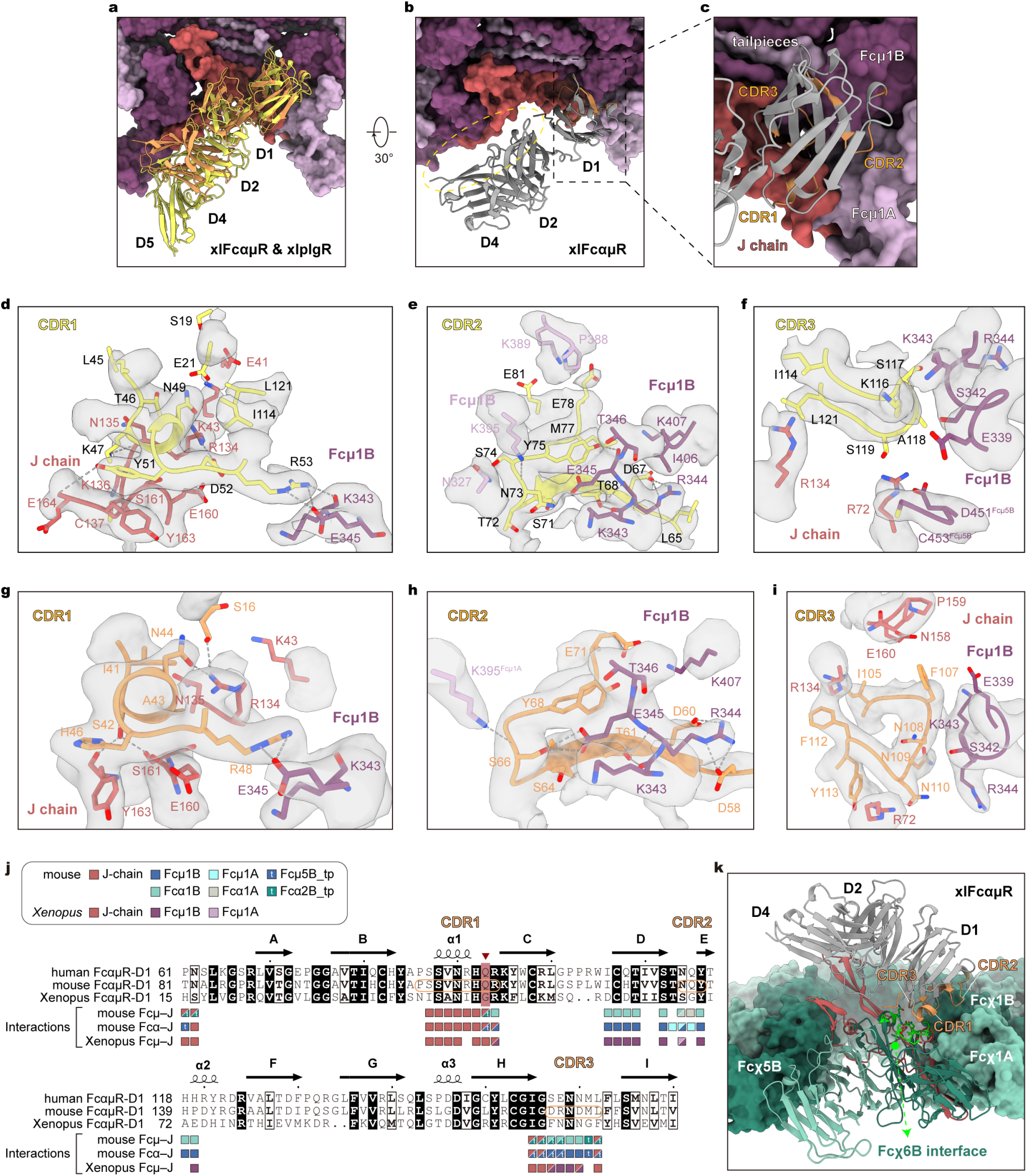
Molecular details of *Xenopus* FcαμR and pIgR binding to pIgs. **a**, Comparison of xlpIgR-ECD and xlFcαμR-ECD binding to xlFcμ–J. FcαμR is shown in sandy brown, while pIgR is shown in yellow. Fcμ–J is displayed as surface. **b,** The xlFcαμR engages Fcμ–J through its D1 domain (black box). Both D2 and D4 domains show no direct interaction with Fcμ–J (black oval). **c,** Binding interface between xlFcαμR and xlFcμ–J. FcαμR-D1 is shown as grey ribbon, with interacting residues highlighted in sandy brown. Fcμ–J is shown as surface. **d–f,** Interactions between xlpIgR-D1 and Fcμ–J at CDR1 (**d**), CDR2 (**e**) and CDR3 (**f**) regions. Corresponding local density maps are shown in grey. Dash lines indicate polar interactions. **g–i,** Interactions between xlFcαμR-D1 and Fcμ–J at CDR1 (**g**), CDR2 (**h**) and CDR3 (**i**) regions. Corresponding local density maps are shown in grey. Dash lines indicate polar interactions. **j,** Sequence alignment of the FcαμR-D1 domain from human, mouse, and *Xenopus laevis*. Residues involved in interactions are evaluated and marked with colored squares. Notably, xlFcαμR exhibits minimal interaction with residues located in the Fcμ1A domain and the tailpiece. The three CDR regions are highlighted with rectangles in the mouse FcαμR sequence. Gly47 (red) in the CDR1 of xlFcαμR increases conformational flexibility of this loop. **k,** Superposition of the xlFcαμR–Fcμ–J complex with the Fcχ hexamer (PDB: 9UO6) reveals compatible docking. The D1 domain occupies a position between two adjacent Fcχ protomers (Fcχ1B and Fcχ6B) with less steric clash, while D2 and D4 make no contact with the Fcχ surface. Although interactions with the J-chain are absent, new potential contacts form with Fcχ6B residues Gln370–Lys379, Pro403, Lys406, and Gln409–Asp411. In the view, the Fcχ molecules are colored in two shades of green. Fcχ1–5 protomers are shown as surfaces and Fcχ6 as ribbon. The Fcχ6B residues potentially involved in xlFcαμR engagement are colored lime.

